# ARID-sf: A physics-informed Deep Learning scoring function to improve Antibody-Antigen docking model ranking

**DOI:** 10.64898/2026.01.20.700530

**Authors:** Ilyas Grandguillaume, Catherine Etchebest, Fernando Luis Barroso da Silva

## Abstract

Accurate prediction of antibody-antigen (Ab-Ag) complexation is crucial for understanding immune responses, diagnostics, and the development of therapeutic antibodies. While molecular docking generates conformations, current scoring functions struggle to identify near-native poses, particularly for Ab-Ag interactions. We present ARID-sf (**A**ntibody-antigen **R**esidue **I**nterface **D**ocking **s**coring **f**unction), which combines classical force field potentials with structural features and protein language model embeddings through a self-attention-based neural network architecture. ARID-sf was trained on >1.5 million docking models and evaluated across four independent test sets comprising 806 cases and 700,000+ docking models. ARID-sf consistently outperforms other functions on increasingly challenging docking scenarios. Critically, ARID-sf maintains performance across diverse sequence identities and increasing conformational change required to reach the bound state, starting with unbound components, demonstrating robust generalization. ARID-sf is parallelizable and can process thousands of docking models per minute, enabling practical application in computational Ab engineering pipelines. The code, trained network, and complete pipeline are freely available at https://github.com/DSIMB/ARID-sf.git.

## Introduction

Antibodies (Abs), also known as immunoglobulins (Igs), are proteins naturally produced by a host’s immune system as a defense mechanism against primarily non-self proteins (Janeway, 2001) but also self ones (Elkon et Casali, 2008). Circulating immunoglobulins of the IgG type are composed of two heavy chains and two light chains (Janeway, 2001). Each heavy chain (H) is composed of three constant and one variable domain, while each light chain (L) is composed of one variable and one constant domain (Janeway, 2001). Each variable domain (H, L) carries three loops (1-3) that are referred to as complementarity-determining regions (CDR), which are often responsible for the antigenic recognition (Reis et al., 2022). The rest of the variable domain, called the framework region, has also been identified to contribute to Ag binding (Masuda et al., 2006). Abs bind their targets, called antigens (Ags), and are then recognized by immune cells such as macrophages (Fridman, 1991), which internalize and degrade the targeted antigen. This binding mechanism involves different components of the Ab, i.e., the constant region that is recognized by the host immune system cells (Fridman, 1991), and the variable region that is responsible for binding the target (Janeway, 2001). In the case of camelidae, the antibodies contain a single domain (sdAb) bearing a single variable heavy chain. They are frequently denominated as “nanobodies”. In this work, we distinguish these two Ab classes that differ by the number of Ab domains: the multi-domain Abs (Variable fragments: Fv, and Antigen binding Fragments: Fab) with at least a light and heavy variable chain, and the nanobodies.

Deciphering the molecular determinants of antibody-antigen (Ab-Ag) recognition is a cornerstone of modern immunology, driving the advancements in vaccine design, diagnostics, and targeted new therapeutics. Classical biophysical methods to study Ab-Ag interaction include crystallography and NMR spectroscopy (Rao et al., 2014), which provide detailed information about the binding site. However, these methods often have important drawbacks; large systems are difficult to study with NMR spectroscopy (Purslow et al., 2020), and optimal crystal conditions must be achieved prior to X-ray diffraction experiments, which requires protein saturation. Moreover, crystallization is often achieved in non-physiological experimental conditions, e.g., pH conditions different than 7 and high salt concentration (Dessau et Modis, 2011). Alternative methods, such as directed mutagenesis (Fontayne et al., 2007), can bring information about residues involved in the interaction.

Alternatively, epitope and paratope locations can be predicted using computational methods based on sequence analysis, structural geometry, or physicochemical signatures such as pKa shifts (Krawczyk et al., 2013; Poveda-Cuevas, Etchebest et Barroso Da Silva, 2020; Clifford *et al.*, 2022; Tubiana, Schneidman-Duhovny et Wolfson, 2022). Therefore, these methods are of major interest as they are less time-consuming, assuming that they are able to provide sufficiently accurate information.

Recently, co-folding methods such as AlphaFold3 or Boltz2 (Abramson et al., 2024; Passaro *et al.*, 2025), leveraging massive multiple sequence alignment and deep learning attention mechanisms, have improved significantly in the task of predicting biomolecular complexes. Although this is also true for Ab-Ag complexes, the prediction success rates for the top-ranked models are still below 50% (Hitawala et Gray, 2025), highlighting the need for further improvements. One of the main aspects preventing reliable predictions of Ab-Ag complexes lies in the difficulty of modelling the Ab CDRs, especially CDRH3, which has an important sequential and structural variability among Abs (Weitzner et Gray, 2017; Fernández-Quintero et al., 2023).

Assuming accurate structural information is available for an Ab and Ag, the most straightforward method to study their complexation is to apply docking approaches. Docking consists of searching for the best relative orientation of two (or more) molecules. The surface of the partners involved is generally sampled in order to find the zones with the best compatibility. Docking includes two principal steps: **a)** model generation and **b)** model selection. Each generated model (pose) is scored with a function that varies significantly depending on the docking tool. (Paggi, Pandit et Dror, 2024).

Multiple docking tools for protein-protein interactions are freely available, such as ClusPro (Brenke et al., 2012), ZDOCK (Pierce, Hourai et Weng, 2011), HADDOCK (Dominguez, Boelens et Bonvin, 2003), RosettaDOCK (Gray et al., 2003), Hex (Ritchie et Kemp, 2000), and ATTRACT (de Vries et al., 2015). Each has its specific scoring scheme and uses different types of algorithms to sample conformations. For instance, ZDOCK and ClusPro employ Fast Fourier Transform algorithms for rigid-body docking, while ATTRACT uses coarse-grained representation with normal modes analysis for accounting flexibility. Hex utilizes spherical polar Fourier correlations for rapid shape matching, and RosettaDOCK combines low-resolution rigid-body search with high-resolution refinement using Monte Carlo optimization and the Rosetta energy function. HADDOCK has been created to incorporate distance restraints initially from NMR experiments to guide docking. Thus, its scoring function includes a useful restraint energy term consistently applied during all docking stages, penalizing geometries that deviate from the expected binding constraints, if the complexation does not satisfy distances between residues. The latest results on Ab-Ag docking present Haddock as one of the best docking tools to generate and rank Ab-Ag docking models with or without any information on interface residues (Ambrosetti 2020). Additionally, the possibility of including specific residues to guide docking in a particular location of the conformational space is valuable to create benchmark sets with an equal number of decoys (false poses, i.e., non-native structure) and high-quality docking models for downstream machine learning applications.

Once the docked models are produced, identification of the best complex is done by selecting the models with the highest score according to the given scoring function (Paggi, Pandit et Dror, 2024). Therefore, the quality of the scoring function is of major importance to accurately predict the 3D structure of a complex.

Historically, docking methods and their scoring schemes have been developed and aimed at general protein-protein interaction (PPi) studies, and have been performing well in this context (Lensink et al., 2020). In contrast, their performance in Ab–Ag prediction has been less satisfactory, both in the generation of high-quality docking models and in their ranking. This limitation also mainly arises from the challenges of modeling the highly variable CDRH3 loop (Harmalkar et al., 2025). Importantly, most docking methods were not originally designed to account for the specific structural and biophysical features of Ab–Ag recognition, which further contributes to the reduced accuracy in this domain. Indeed, Abs paratopes have been extensively studied, and specificities such as the enrichment of Tyrosine (a titratable amino acid) and other sequential patterns (Akbar et al., 2021; Reis et al., 2022) have led to the identification of clear Ab interface markers. Inversely, the prediction of epitope properties is still a major challenge (Van Regenmortel, 2014; Cia, Pucci et Rooman, 2023). Thus, Ab-Ag interactions are challenging to predict, and there are improvements to be made both in the generation and ranking of Ab-Ag docking models to improve the overall reliability of computational methods on the subject.

Recent efforts to develop Ab-specific scoring functions have shown promise but remain limited in scope. DLAB (Schneider et al., 2022) combines ZDOCK (Pierce, Hourai et Weng, 2011) and convolutional neural networks with 3D grid representations to improve pose ranking and virtual screening, showing particular strength when working with modeled Ab structures. ClusPro (Desta et al., 2023) incorporates Ab-specific parameters in its scoring scheme. These two scoring functions are tied to their docking software. The CSM-AB (Myung, Pires & Ascher, 2022) scoring function employs graph-based representations to predict binding affinity from structural features, demonstrating improvements over general-purpose methods. AbEpiScore predicts both model quality and epitope location using AlphaFold-based approaches and inverse folding techniques (Clifford et al., 2025). Despite not being tied to any docking software, these two scoring functions are computationally expensive when used for high-throughput studies of tens of thousands of docking models, as the scoring takes significant time compared to the production of the docking model (by a physics-based docking tool such as ZDOCK or HADDOCK).

To the best of our knowledge, the field lacks a highly specialized function able to accurately rank docking models from any source involving Fab, Fv, or sdAbs that is applicable for high-throughput docking studies involving tens of thousands of models while maintaining consistent performance across different docking protocols and structural perturbations.

### Approach/Strategy

In the present work, we develop a new scoring function, **ARID-sf** (**A**ntibody-antigen **R**esidue **I**nterface **D**ocking **s**coring **f**unction), which combines classical physical force field potentials with structural features and protein language model embeddings through a self-attention-based neural network architecture. Rather than developing a new docking method, we created a new specialized scoring function designed to evaluate and rank Ab-Ag docking models (poses) generated by existing tools. We selected HADDOCK3 (Giulini et al., 2025) as the associated tool for model generation due to its modular design, restraint options, and proven effectiveness for Ab-Ag docking (Ambrosetti et al., 2020). ARID-sf combines physicochemical, structural, and sequence features through an attention-based neural network.

Our central hypothesis is that physics-based scoring functions accurately capture general physical interactions but might miss unique characteristics of Ab-Ag recognition. Specifically, incorporating physical potentials of interactions as features allows the neural network to leverage established biophysical principles rather than learning them *de novo* from the training data. Additionally, it enables the function to contextualise the physical contribution within the local interface environment. The additional context we provide consists of ESM embeddings, i.e., numerical representations of protein sequences learned by large-scale language models (Hayes et al., 2025), interface volumic properties (see the method section), as well as atom-atom or residue-residue contacts, all features that are useful to predict interaction properties such as binding affinity (Vangone et Bonvin, 2015; Miller et al., 2023). ARID-sf is able to rerank Ab-Ag docking models with improved precision across all docking stages while maintaining computational efficiency suitable for large-scale screening.

## Material and Methods

In the present section, the main protocols, tools, and metrics used throughout the results are described. The complete details for reproducibility, including parameters, hyperparameters, architecture details, configuration files, and additional results, can be found in the supplementary material (SM), with explicit cross-references between theSM sections and the main paper.

### Dataset Creation for Training

In this work, we refer to experimental complex (crystallographic, NMR, or Cryo-EM structures) as “cases” and their components (Ag and Ab) as the native bound structures. The initial dataset was derived from 2,151 Ab-Ag complexes extracted from the ANABAG dataset (Grandguillaume, Barroso Da Silva & Etchebest, 2025). To minimize sequence redundancy and prevent data leakage during training, we applied sequence identity thresholds separately for antigens and antibodies (see **SM**: clustering parameters). This filtering resulted in a curated dataset of 1,046 non-redundant cases. Each structure file was preprocessed such that antigen atoms were placed first, followed by antibody atoms, with continuous renumbering from 1 to ensure consistency across all cases.

The dataset was partitioned into three non-overlapping sets based on 20% antigen sequence identity clustering, yielding 263 distinct sequence clusters. The training set contains 997 cases from 230 clusters, the validation set includes 34 cases from 18 clusters, and the internal (i.e., set created from the 1,046 cases) test set comprises 15 cases from 15 clusters. This stringent partitioning ensures that no antigen sequence in the test or validation set shares more than 20% identity with any sequence in the training set, providing robust evaluation of generalization performance.

### Docking Model Generation

Docking models serve as the structural basis for training ARID-sf, requiring diverse quality distributions from incorrect to high-quality predictions (Figure 1, Top panel). Model quality was assessed by comparing interfaces against reference complexes using the caprieval module from HADDOCK, with stratification into four CAPRI quality categories based on DockQ scores (Basu et Wallner, 2016; Collins et al., 2024).

**Figure 1.**
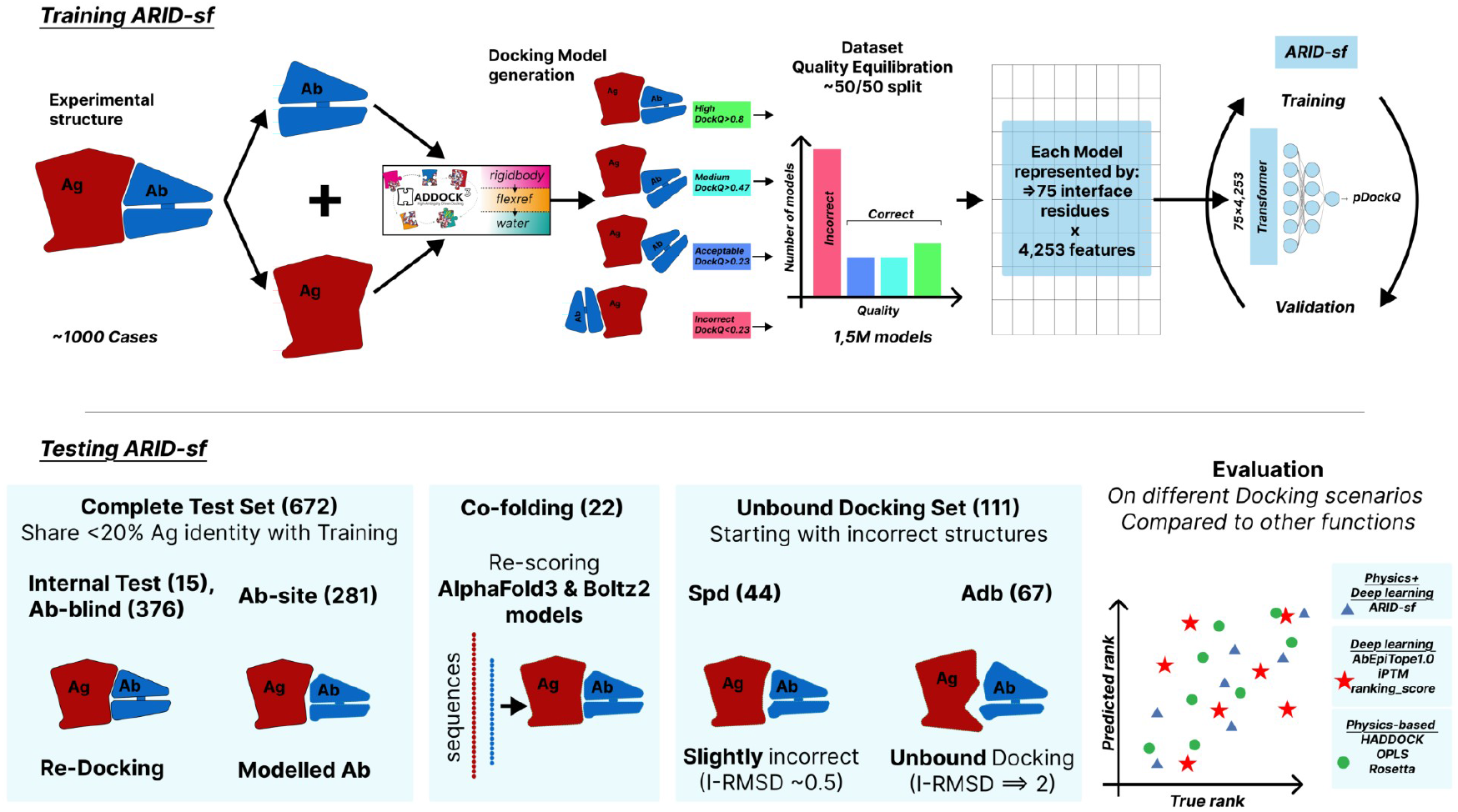
Workflow describing how ARID-sf was trained (top) and evaluated (bottom). ***(Top)*** Training: ARID-sf was trained using docking models generated with HADDOCK3 and separated components of Ab-Ag experimental structures. The three HADDOCK modules *(rigidbody, flexref, and water)* have been used to generate the docking models. The entire dataset has been equilibrated in terms of docking model quality, resulting in a ~50% split between correct and incorrect models. Each model is described by a feature matrix of up to 75 residues in length and 4,253 residues in width. ARID-sf was trained and validated on this feature representation to predict the *DockQ* score of the corresponding model, using a transformer architecture. ***(Bottom)*** Evaluation: ARID-sf was then evaluated on several datasets of docking models: the complete test set aims at evaluating ARID-sf on structures having no antigen homology with the training set, and represents the baseline evaluation. ARID-sf was also evaluated on models produced by co-folding methods, to assess the function biases towards HADDOCK generated models. Finally, ARID-sf was evaluated on unbound docking, with the *Spd* set, representing slight perturbations of the interfaces, and the *Adb* set, including unbound starting structures requiring substantial changes to reach the correct state. The evaluations are based on ranking and classifications tasks, and ARID-sf is systematically compared to other physics or deep learning based, state of the art functions.

We employed HADDOCK3 (Giulini et al., 2025) with a fully guided protocol to generate training models. For each of the 1,046 training-eval-test cases, we generated 3,000 rigid-body docking models by **i)** identifying interface residues from reference complexes (see SM: interface definition), **ii)** creating restraint files containing interface residue lists, and **iii)** executing *rigid-body* docking guided to the interface region (see SM: HADDOCK protocol details). This yielded approximately 3 million models. To balance quality distribution during training, we randomly selected 1,000 models per case with target proportions of approximately 500 incorrect, 125 acceptable, 125 medium, and 150 high-quality models, resulting in approximately 1 million rigid docking models and a 50/50 split between correct and incorrect models. Then, to capture interfacial adaptations of correct (but mostly) incorrect poses, we applied flexible refinement protocols: **i)** flexibility refinement using the HADDOCK “flexref” module, and **ii)** water refinement (see SM: refinement protocols). The “flexref” and water structures are considered separately. However, to diminish the computational costs, we only performed refinement on a subset of 479 cases representing half of the total pool of cases. From 479 cases, we selected 125 models from each quality category (500 total models). This expanded the dataset to 1,500 models for these 479 cases. The complete training dataset comprised 1,544,000 models (1,065,000 rigid and 479,000 flexible refined models) spanning the full quality distribution required for training (see Table 1).

**Table 1:** Composition of each training or test set. The number of Cases is reported with the total number of models obtained with the HADDOCK “rigidbody” module or “flexref/mdref” module (Flexible Models). The type of experiment is also reported, either re-docking for docking separated components of solved complexes, or unbound docking where one or all components deviate from the native complexed structure.

|  | Cases | Rigid Body Models | Flexible Models | Type |
| --- | --- | --- | --- | --- |
| Train | 997 | 1,015,118 | 456,607 | Re-docking |
| Evaluation | 34 | 34,612 | 15,538 | Re-docking |
| Test (internal) | 15 | 15,270 | 6,855 | Re-docking |
| Complete Test Set | 672 | 15,270 | 171,105 | Re-docking/<br>Unbound docking |
| Spd | 44 | 132,000 | - | Unbound docking |
| Adb | 67 | 268,000 | - | Unbound docking |

Subsequently, each interface is represented by several features that are processed by the ARID-sf deep learning algorithm. The targeted function is the DockQ score (Figure 1, Top panel, right side, for all the details see “Features calculated by ARID-sf” section and “Predicted Properties”)

### Dataset Creation for Testing

Beyond the internal test set, we assembled three additional test sets to comprehensively evaluate ARID-sf performance across diverse scenarios and docking protocols (Figure 1).

#### Complete Test Set (Ab-blind and Ab-site)

We incorporated docked structures from the Ab-Set. Ab-Set is a comprehensive dataset recently released by Almeida et al. (Almeida et al., 2025), which employed two generation strategies: **i)** a blind protocol consisting of re-docking without interface restraints (Ab-blind set), and **ii)** a targeted protocol guided towards the interface using ABodyBuilder2-predicted antibody structures (Ab-site set) (Leem et al., 2016). From these datasets, we selected 376 cases from Ab-blind and 281 cases from Ab-site, excluding any cases sharing more than 20% Ag or 95% Ab sequence identity with the train set cases. These models (384,496 for Ab-blind, 323,998 for Ab-site, and 27,419 for the 15 cases internal test) were combined to form the primary test set for evaluation.

#### Slight Perturbation Docking Set (Spd)

To evaluate scoring function robustness under conditions where epitope and paratope conformations deviate from the reference bound complex, we created a specialized test set using the Ab-Set. We selected 100 representative cases with low sequence identity to the training set (<20% for Ag, <80% for Ab). Rather than using native Ab and Ag components, we extracted components from previously generated docking models that exhibited conformational shifts relative to native structures (paratope-paratope and epitope-epitope Root Mean Squared Deviation (RMSD) fluctuating around 0.5 Å). These perturbed components were then subjected to semi-blind rigid-body docking (see SM: restraint protocol), generating 3,000 models per case. Among these 100 cases, 44 yielded acceptable to high-quality models suitable for evaluation (see SM: detailed composition).

#### ABAG-Docking Benchmark Set (Adb)

To assess ARID-sf performance on complexes with substantial interfacial conformational variability, we utilized 67 Ab-Ag cases from the ABAG-docking benchmark (Zhao et al., 2024). This benchmark provides both bound and unbound protein structures, enabling evaluation of the capability of the scoring function to rank correct poses starting from structurally incorrect conformations. We generated 4,000 rigid-body decoys per case using HADDOCK3 following a semi-blind protocol (see SM: protocol details), with 39 of 67 cases producing at least one acceptable-quality model. Structural quality metrics were calculated against bound reference structures (see SM: metric details).

#### AlphaFold3 and Boltz2 Test Sets

To evaluate ARID-sf on models generated by deep learning inference frameworks, we selected 16 cases absent from both AlphaFold3 and Boltz2 (Abramson et al., 2024; Passaro et al., 2025) training data, plus 6 cases absent only from AlphaFold3 training data (see SM: complete case list). For both AlphaFold3 and Boltz2, 25 models were generated per case (5 seeds × 5 models), and Boltz2 was configured to replicate AlphaFold3’s default parameters (see SM: specific parameters). Due to computational cost, the number of models was limited to 25, although the results from Boltz2 were obtained on average with a threefold speedup compared to AlphaFold3.

### Features Calculated by ARID-sf

ARID-sf employs a two-stage process: interface feature calculation followed by neural network inference. Interface residues are identified using a 5 Å heavy atom distance cutoff between Ab and Ag components, a standard criterion in the field (Collins et al., 2024). Each interface residue serves as a central point (named “central residue”, see Figure 2) for multi-scale feature encoding, integrating evolutionary information, physical interactions, and geometric descriptors into a 4,253-dimensional feature vector.

**Figure 2:**
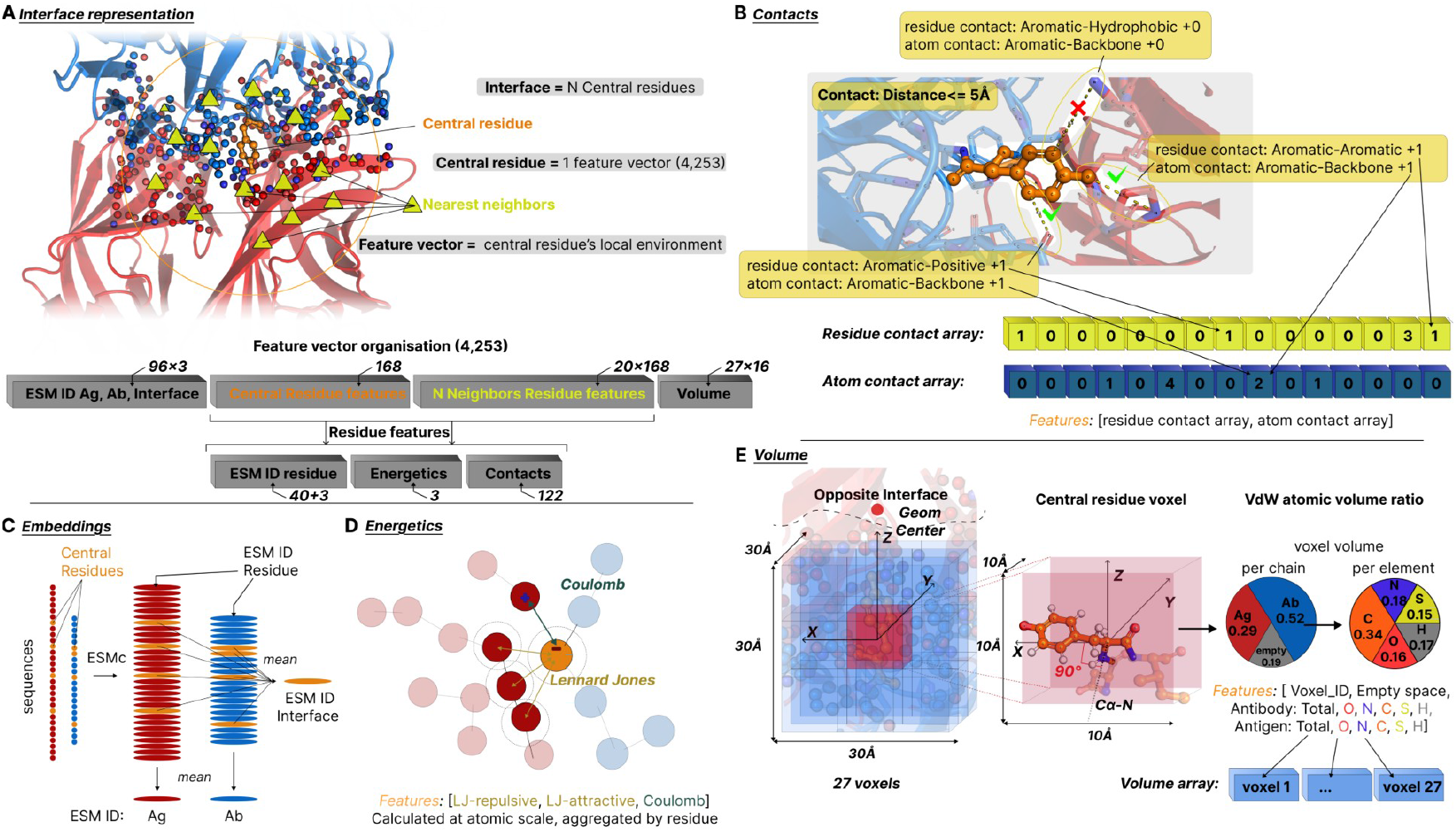
Features utilized in ARID-sf. ***A)*** Interface representation of the docking model to be scored. Each amino acid at the interface will constitute a central residue, associated with a feature vector of size 4,253 describing its local environment. Up to 75 central residues are described for a given model. The feature vector consists of four main parts: ESM identifiers (ID) (see C), Central and Neighbors residue features, and finally the volume description (see E). The central and neighbor residue features are organized the same way: an ESM ID encoding the corresponding residue, energetics (see D), and contacts (see B). ***B)*** Contact description is made at the atomic and residue level. The contacts array is filled with integers, counting the total number of contacts of one particular combination, for instance, aromatic-aromatic. ***C)*** ESMc embeddings are extracted from the Ag and Ab sequences. Embeddings are aggregated by mean pooling for the Ag and Ab, creating their ESM ID. Interface residues (orange) are also aggregated to create the ESM Interface ID. Individual amino acid embeddings are used as residue ID in the residue features (see A). ***D)*** Energetics are calculated using the united atom representation of the structure, and aggregated by residue. ***E)*** Volume properties are calculated by placing a cube centered around the central residue. The cube is centered consistently using the backbone atoms and the opposite interface geometrical center. Each voxel contains atoms of the Ag or Ab chains. In each voxel, the fraction of occupied volume of each element and its associated component (Ag or Ab) is reported using the atom VdW radius. Voxel features are then stacked in the same order to create the volume array (see A).

#### Feature dimensions

The input representation for each docking model comprises a matrix with dimensions equal to the number of central interface residues (up to 75) in length and 4,253 features in width (Figure 1 top). Residues are ordered by decreasing intermolecular contacts, ensuring the 75 most connected residues are prioritized when interfaces exceed 75 residues. This threshold captures 95% of observed interface sizes in the training data, with an average interface of 45 residues. The creation and decomposition of the 4,253 feature vector is shown in Figure 2.

#### Antigen, Antibody, Interface, and residue identifiers (ID)

Identifiers are used to signify to the deep learning model the sequence information of the complex. Residue-specific profiles are extracted using the ESM Cambrian (ESM-C) model (Hayes et al., 2025). Residue embeddings are reduced in length (see SM: ESM extraction details) and used as residue ID in residue feature arrays (Figure 2 A, C). The Ag, Ab, and Interface ESM IDs are all obtained by aggregating residue embeddings for the entire Ag, Ab, and Interface sequence, respectively.

#### Physical Interactions

Non-bonded interactions are calculated using OPLS-UA force field parameters matching those employed in HADDOCK docking calculations (Jorgensen et Tirado-Rives, 1988; Vangone et al., 2017). The Lennard-Jones 6-12 potential is explicitly decomposed into separate repulsive and attractive components, while Coulombic interactions are computed with appropriate cutoff functions (See Figure 2 D and SM: detailed parameters and validation). This separation enables the neural network to learn optimal weighting for each physical contribution as a function of the local context.

#### Contact Maps

Residue-residue and atom-atom interactions within 5 Å are captured and categorized by physicochemical properties (Figure 2B). Residues are classified into five groups: hydrophobic, polar, aromatic, negatively charged, and positively charged (see SM: complete classification). Similarly, atoms are categorized as backbone, hydrophobic, polar, aromatic, negatively charged, and positively charged (see SM: atom categorization details). Multiple contacts between the atoms of the residue are only counted once at the residue contact level.

#### Volumetric Features

A 3×3×3 voxel grid (10 Å per voxel) is constructed around each central residue Cα atom (Figure 2E). The cube size of 30 Å is identical to what has been used for DeepRank, a scoring function for non-immune interactions and crystal artifact identification (Renaud et al., 2021). The grid orientation is established by i) positioning the center at the Cα atom, ii) aligning the Z-axis from Cα toward the opposite interface center, iii) defining the X-axis perpendicular to both Z and the Cα-N backbone vector, iv) completing the orthonormal system with the Y-axis, and v) calculating atomic volumes within each voxel using radii. This representation encodes spatial occupancy of the Ag and Ab atoms as well as empty space fractions, providing a description of the local environment as a proxy for solvent accessibility (see SM: detailed calculation method).

### Predicted Property

ARID-sf predicts the DockQ score (pDockQ) of each docking model, providing a single quality metric that integrates multiple structural similarity measures. DockQ ranges from 0 to 1, with values corresponding to the standard CAPRI quality categories: Incorrect (<0.23), Acceptable (0.23–0.47), Medium (0.47–0.80), and High (>0.80) (Basu et Wallner, 2016; Collins et al., 2024). This metric serves as the target for training the neural network and enables direct comparison of predicted quality against ground-truth structural accuracy calculated from reference complexes.

### Training

The neural network architecture is based on transformer encoders, designed to process variable-length interfaces and predict docking model quality (Figure 2D). The high-dimensional input features (4,253) first undergo linear transformation to a 256-dimensional representation space with layer normalization and dropout regularization, enabling both compact representation learning and data normalization for subsequent transformer processing. For more details about the architecture, see (SM: Neural Network Architecture and Training). The function used for training is a root mean square error (RMSE) loss function optimized using the Adam optimizer (see SM: Neural Network Architecture and Training).

### Evaluation Metrics

To evaluate the scoring functions, we used the Success Rate, Hit Rate, and Minimum Misclassification Rate. The Success Rate corresponds to the common definition used to evaluate other scoring functions in the literature (Xue et al., 2014; Renaud et al., 2021), and represents the main evaluation metric used in the results section. The Hit Rate is defined in this work to account for the number of correct and incorrect models in the top predictions. The Minimum Misclassification Rate is a new evaluation tool that helps identify the optimal score threshold above which a model should be correct (see example below).

Success rates (SR) are calculated as the cumulative fraction of cases where at least one correct model is found following the in the top N using:

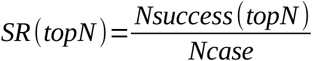

Where *Nsuccess(topN)* is the number of cases where at least one correct model is found in the *topN* models, and *Ncase* is the total number of cases. For instance, a SR of 10% for topN = 10 signifies that for 10 percent of cases (different complexes) at least one correct model is found in the top 10 according to the tested scoring function.

Hit Rate (HR) within the top-N ranked models quantifies the ratio of correct poses (acceptable quality and above) in the top N, regardless of the number of correct cases. For a given top-N selection, the HR is calculated as:

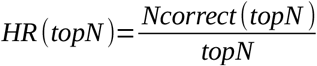

where *Ncorrect(topN)* represents the number of correct poses among the *topN* selected models. HR converges toward the overall rate of correct poses in the ensemble as *topN* approaches the total number of generated models. For instance, an HR of 10% for topN = 10 signifies that 10 percent of models are correct in the top 10 selected models. The theoretical best possible performance is calculated as:

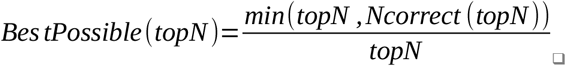

The minimum misclassification rate (MCR) evaluates the ability of the scoring scheme to distinguish correct from incorrect poses by identifying the optimal classification threshold. MCR is computed as:

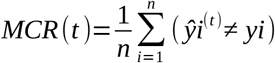

where *t* is the classification threshold, *ŷ* is the predicted quality at threshold *t, yi* is the true quality, and *n* is the number of evaluated models. Lower MCR values indicate superior discriminative capacity. For instance, an MCR of 10% with t = 0.5 signifies that the best achievable classification accuracy for the scoring function is 90% by using 0.5 as the score threshold value, assuming correct prediction above 0.5. The threshold *t* yields the lowest MCR values for each set.

### Comparison with Other Scoring Functions

We evaluated ARID-sf against several established scoring schemes developed prior to this work. The HADDOCK scoring scheme was assessed without the restraints penalty term to enable fair comparison, as this term requires prior interface knowledge. We systematically applied scoring functions with weights corresponding to the docking protocol as described by Vangone et al. (Vangone et al., 2017). The HADDOCK score comprises electrostatic and potential terms, an empirical desolvation term, and buried surface area contributions (see SM: protocol-specific weights).

HADDOCK3 reports and electrostatic potentials calculated using the OPLS united atom force field. The “OPLS scores” represent unweighted interaction energies (sum of and electrostatic contributions). Both HADDOCK and OPLS scores are transformed into ranks, with the best model (minimum energy) ranked first, enabling comparison with other scoring functions.

AbEpiScore-1.0 and AbEpiTarget-1.0 were calculated using the AbEpiTope-1.0 software (Clifford et al., 2025) for each of the 3,000 models in the Spd set. The calculation required approximately 3 days for 132,000 models using the same hardware employed for ARID-sf training, compared to approximately 1 hour for ARID-sf scoring.

Rosetta scores were obtained by running the interface analyser using the REF2015 scoring function (Alford et al., 2017) on each model of the Spd set. Default parameters were used, and the total_score is used as the ranking score.

Both Boltz2 and AlphaFold3 (Abramson et al., 2024; Passaro et al., 2025) were evaluated to assess ARID-sf performance on models generated outside the HADDOCK framework. We retrieved the ipTM (Interface Predicted Template Modeling score), pTM (global Predicted Template Modeling score), and ranking_score (termed “confidence_score” in Boltz2) for each generated structure. Since pTM consistently underperformed the other scores on the evaluated tasks, we focused comparisons on ipTM and ranking_score. Structures from both platforms were converted to PDB format, processed through the HADDOCK3 topoaa module for topology compatibility, and subsequently scored using ARID-sf (see SM: detailed processing pipeline).

## Results

ARID-sf algorithm predicts DockQ scores based on per-residue environmental features. While designed as a regression model, it can function as a classifier with appropriate thresholds. We compared ARID-sf against two physics-based functions: the HADDOCK score (weighted OPLS potential with desolvation term and buried surface area depending on the protocol) and the OPLS score (unweighted and electrostatic terms). The performances were evaluated on four datasets: the complete test, Spd, Adb, and a dataset of Boltz2 and AlphaFold3 generated complexes.

The scoring functions’ performances are evaluated through the classical Success Rate (SR), the Hit Rate (HR), and the Minimum MisClassification rate (MCR) (see the Methods section). The SR performances of ARID-sf, the HADDOCK scoring function, and OPLS score are summarized in Table 1. Each set evaluation is presented in its own result section.

### Performances of ARID-sf to rank and classify docking structural models

The complete test set represents the baseline evaluation of ARID-sf on an ensemble of 672 Ab-Ag cases, obtained via different docking protocols, and through which ARID-sf is compared with the HADDOCK and OPLS scores. SR of ARID-sf are higher than HADDOCK and OPLS scores, for this test set, signifying that ARID-sf provides a better Ab-Ag model ranking compared to the other functions (see Figure 3A, Table 2).

**Table 2:** Success rates for scoring functions on the Test, Spd, and Adb set. Success rate represents the fraction of cases with at least one correct model identified within the top 1 or 10 ranked positions (shown in Figure 3D, Figure 4, and Figure 6A). Values are mean ± standard deviation. Statistical significance (α = 0.05, two-sample t-test): *h indicates p < 0.05 versus HADDOCK values; *o indicates p < 0.05 versus OPLS values; *a indicates p < 0.05 versus AbEpiScore-1.0 values; *r indicates p < 0.05 versus Rosetta total_score values. A scoring function appearing with * has a significantly lower Success rate than the scoring function whose column it appears in.

| Scoring | ARID-sf |  | HADDOCK score |  | OPLS score |  |
| --- | --- | --- | --- | --- | --- | --- |
| Top | 1 | 10 | 1 | 10 | 1 | 10 |
| Test set (672) | 0.46± 0.49<br>(*o, *h) | 0.81± 0.38<br>(*o, *h) | 0.39± 0.48 | 0.66± 0.47 | 0.37± 0.48 | 0.61± 0.48 |
| Spd set (44) | 0.93± 0.25<br>(*o, *h, *a, *r) | 0.93± 0.25<br>(*h, *a, *r) | 0.40± 0.49<br>(*a, *r) | 0.47± 0.50<br>(*a) | 0.70± 0.46<br>(*h, *a, *r) | 0.90± 0.29<br>(*h, *a, *r) |
| Adb set (67) | 0.15± 0.36<br>(*o) | 0.20± 0.40 | 0.12± 0.33<br>(*o) | 0.17± 0.38 | 0.0± 0.0 | 0.10± 0.30 |

**Figure 3:**
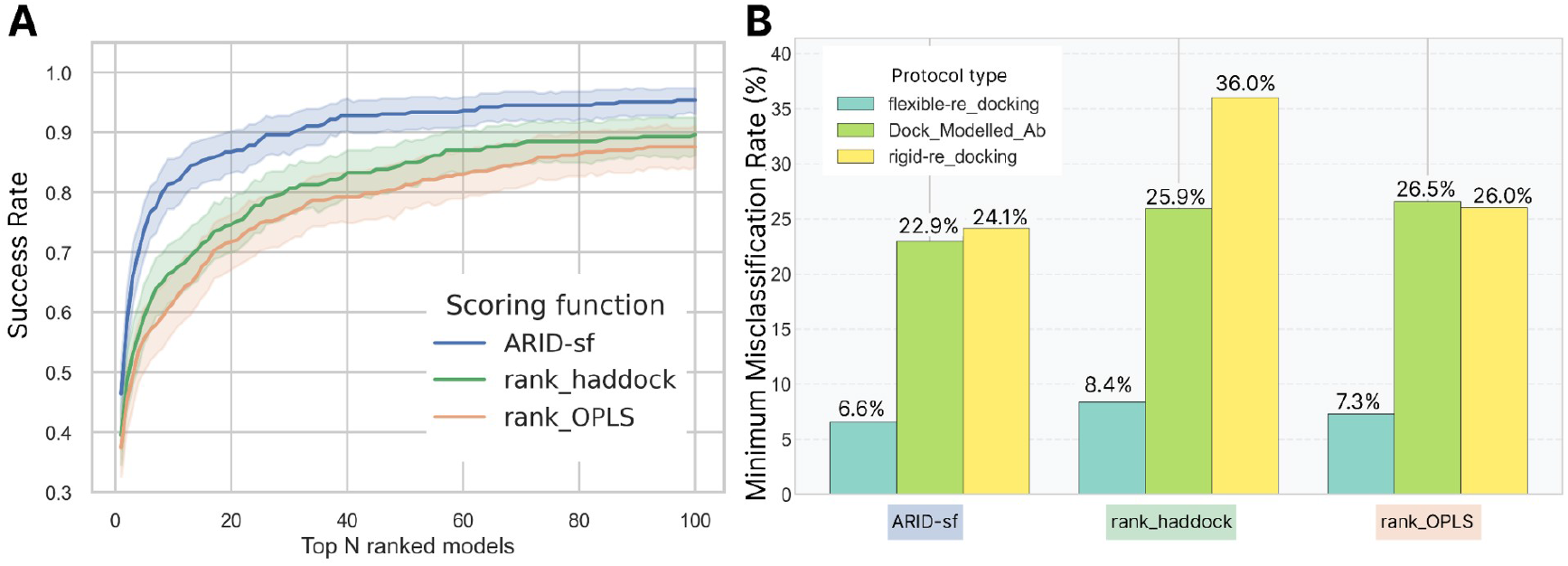
Performance comparison of scoring function. ***A)*** SR curves showing the fraction of cases with at least one correct model in the top-N selections. Shaded areas represent 95% confidence intervals. ***B)*** Minimum misclassification rates for distinguishing correct from incorrect models, stratified by type of protocol used to obtain the docking models (flexible or rigid re-docking and docking of modelled Ab). Lower values indicate better classification accuracy at optimal thresholds.

For top-1 and top-10 ranking (Table 2), ARID-sf achieves SR of 46% and 81%, compared to HADDOCK 39% and 66%. OPLS performs the worst with only 37% and 61% SR. ARID-sf SR performances represent a significant increase compared to both HADDOCK and OPLS, and outperform the physics-based function in ranking.

An important aspect of scoring functions lies in their capacity to distinguish correct from incorrect models at a given threshold score. In the present comparison, all scoring functions show comparable classification ability provided that optimal thresholds are applied (Figure 3B). However, the MCRs are slightly lower for ARID-sf compared to OPLS and HADDOCK, demonstrating that the trained scoring function widens the score gap between correct and incorrect models. As the MCRs have distinct average values for different docking protocols, we differentiated them in this section. ARID-sf achieves minimum misclassification rates of 6.6%, 22.9% and 24.1% for models in the test set coming from flexible-re_docking, modelled-Ab, and rigid-re_docking, respectively (see Methods: Dataset Creation for Testing). The optimal thresholds (*t* see MCR definition in the Materials and Methods section) for ARID-sf are 0.57, 0.23, and 0.34, respectively.

The fact that for different docking experiments, the MCR threshold yielding the best correct-incorrect classification is different highlights an instability of the score. Ideally, the score threshold would be universal for all experiments and cases, allowing one to give the ARID-sf score value above which one should or should not consider a docking model.

Interestingly, the optimal threshold for the modelled-Ab subset corresponds to 0.23, which is the DockQ CAPRI threshold to consider correct models (see Methods, predicted property), highlighting the adequacy between the predicted ARID-sf DockQ and true model quality. As the three scoring schemes share a forcefield foundation (in OPLS united atom, see Methods and SM), the MCR improvements presented in Figure 3 suggest that adding contextual information in addition to the energetic description helps the function in classifying docking results.

### ARID-sf re-scoring of AF3 and Boltz2 generated complexes

In this section, we examine the capacity of ARID-sf to compete with the scoring functions used in deep learning 3D model prediction, i.e., whether ARID-sf achieves better performance in ranking deep learning 3D models compared to the co-folding methods’ ranking schemes. In this aim, we chose two methods that are considered among the best ones for the prediction of complexes, *i*.*e*., AlphaFold3 and Boltz2. ARID-sf scores were calculated for each of the resulting models and compared to the co-folding methods iPTM and ranking_score (see Methods Comparison with Other Scoring Functions). 22 structures have been selected based on their publication date in the PDB databank, and we generated 25 models of the complexes with AF3 and Boltz2, using the corresponding Ab and Ag sequences. For more details, see the Method and SM sections. Among the 22 cases, AlphaFold3 and Boltz2 predicted correct models for 13 cases and 11 cases, respectively, where 7 cases and 8 cases include both correct and incorrect models. To evaluate the scoring functions’ ranking ability, we considered only the cases where both correct and incorrect models were predicted. In total, 175 AF3 models and 200 Boltz2 models were considered in the subsequent SR analysis.

As denoted by the SR curves in Figure 4A, the performances of ARID-sf, although not significantly superior, are better than the ranking_score and iPTM native to AlphaFold3 or Boltz2, for models generated by these co-folding methods.

**Figure 4:**
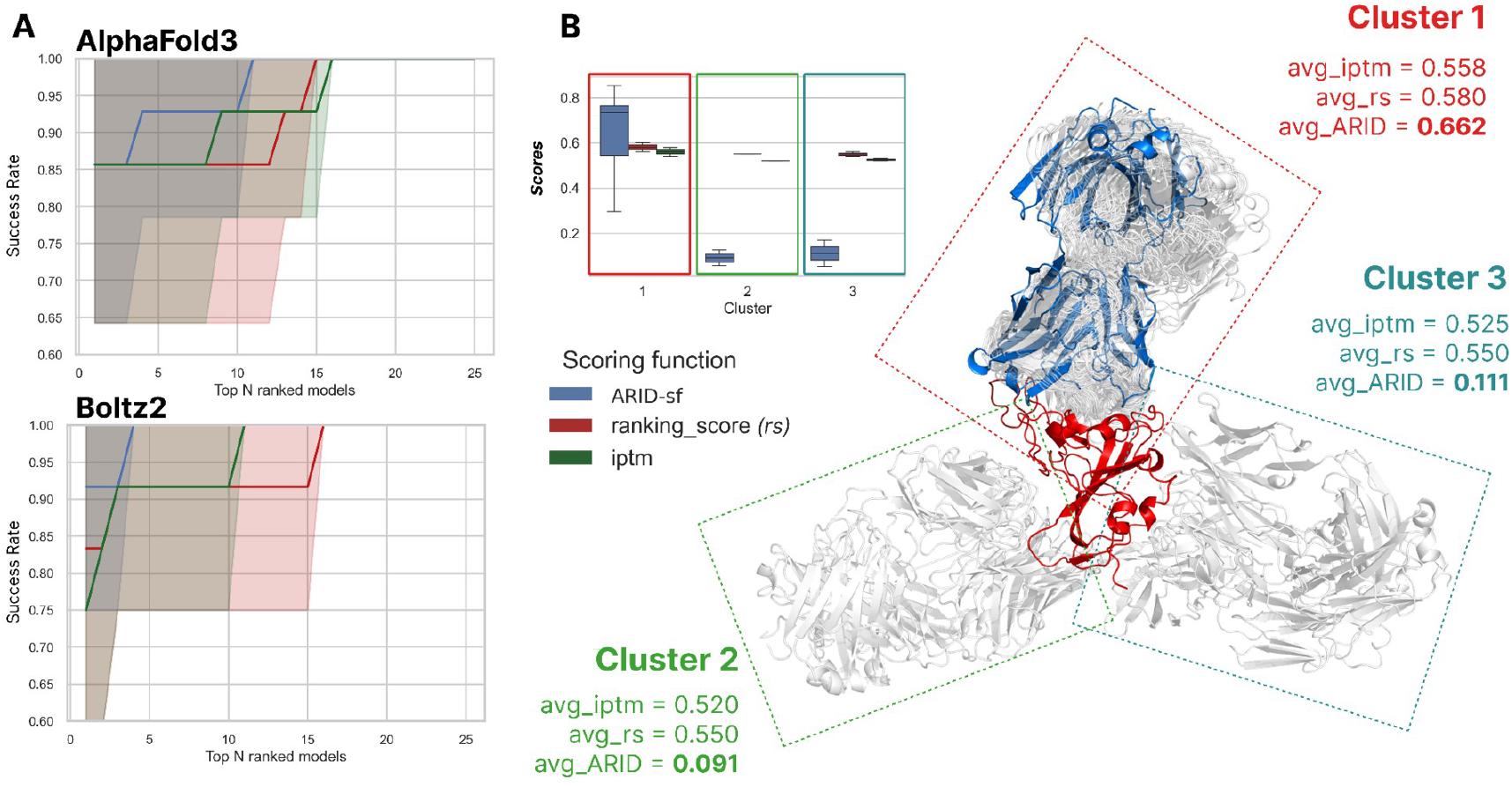
Comparison of ARID-sf with Boltz2 and AlphaFold3 scoring functions. ***A)*** The figure shows the SR for three scoring functions (ARID-sf in blue, the IPTM score in green, and the ranking score in red). The top and bottom plots present results relative to the AlphaFold3 and Boltz2 produced models and scores, respectively. ***B)*** A specific example where all predicted models are Incorrect, and three clusters have been predicted by AlphaFold3. The figure presents the score distributions per cluster in a boxplot, and the average scores are written. The three clusters are outlined by three colors: Cluster 1 in red, Cluster 2 in green, and Cluster 3 in turquoise. The reference Complex is also shown in red for the Ag and blue for the Ab.

Overall, the performances of ARID-sf compared to scores derived from inference models (IPTM and ranking_score) are similar, with slight improvements brought by ARID-sf. The IPTM score has been more reliable on 2 distinct cases compared to the ranking_score, for Boltz2 and AlphaFold3. Because of the significant computational time taken by these inference models to predict the structure and complex from the sequence, a proper comparison would at least require conformational sampling; however, this demonstrates that ARID-sf can be used to score structures that have been produced by other means than HADDOCK.

A marked difference between these inference models and a scoring function such as ARID-sf is the diversity of scores given to a “family” of poses. During docking or inference, it often happens that distinct clusters of highly similar poses are sampled. Depending on the approach, one might score the poses as a function of the cluster population, by the individual quality of each model, or by the average quality of the models in each cluster. Usually, the pose selection is easier if the different families have distinct scores or distinct structural characteristics. We selected one of the cases where only Incorrect models were produced by AlphaFold3 to highlight an important difference between ARID-sf and the IPTM or ranking_score. For this case, 25 models have been produced, and three clusters of structures have been sampled and are presented in Figure 4B. Cluster 1 has the majority of structures (21) and corresponds to the correct orientation of the Ab-epitope association. Clusters 2 and 3 each contain 2 models, with very distinct orientations compared to Cluster 1. Yet, despite these differences, all structures are predicted with very similar iPTM and ranking scores, intra and inter Clusters. In contrast, as displayed in Figure 4B, the scores given by ARID-sf vary substantially between Cluster 2,3, and Cluster 1, and also vary for different models in Cluster 1. In this case, ARID-sf enables easier selection of the correct Ab-Ag orientation by having a clear score separation between the three families of poses. This denotes a sensitivity of the function to more subtle changes in the complex conformation, a property that arises both from the physical description and chemical context used for training ARID-sf.

### Sensibility of the scoring function to interface structural modifications

In the previous section, we showed that ARID-sf has an improved sensitivity for scoring Ab-Ag models compared to pure physics-based and deep learning based scoring functions. How this sensitivity is challenged by small changes in the interface is the question addressed in the present section. The input Ab and Ag spans slight structural deviations, interface RMSDs ranging from 0.35 to 1 Å, with 50% below 0.55 Å (Figure SM S4). Global structure RMSDs cluster slightly below the interface average around 0.5 Å, confirming that perturbations primarily affect binding interfaces (Figure SM S4). Moreover, we add two functions to the comparison pool, the Rosetta total_score, a state-of-the-art scoring function utilized for docking and engineering (Alford et al., 2017), and the AbEpiTope-1.,0 a novel deep learning based scoring function able to distinguish true from false Ab-Ag association and rank docking models (Clifford et al., 2025).

All previously evaluated scoring functions maintain their ranking ability despite the structural perturbations (Figure 5A). ARID-sf and OPLS_rank achieve the highest HR rates with ~80% correct models in the top 10. Interface RMSD shows minimal correlation with performance for small perturbations (Figure SM S4). The Rosetta total_score performs worse than the haddock_rank, with ~44% SR in the top 10 against ~56% SR. Despite its sophisticated AlphaFold-based architecture and inverse folding approach, AbEpiScore-1.0 underperforms on this dataset. While it assigns appropriate average scores to different quality tiers and improves on epitope identification tasks (see SM Figure S2), it struggles with false positive discrimination.

**Figure 5:**
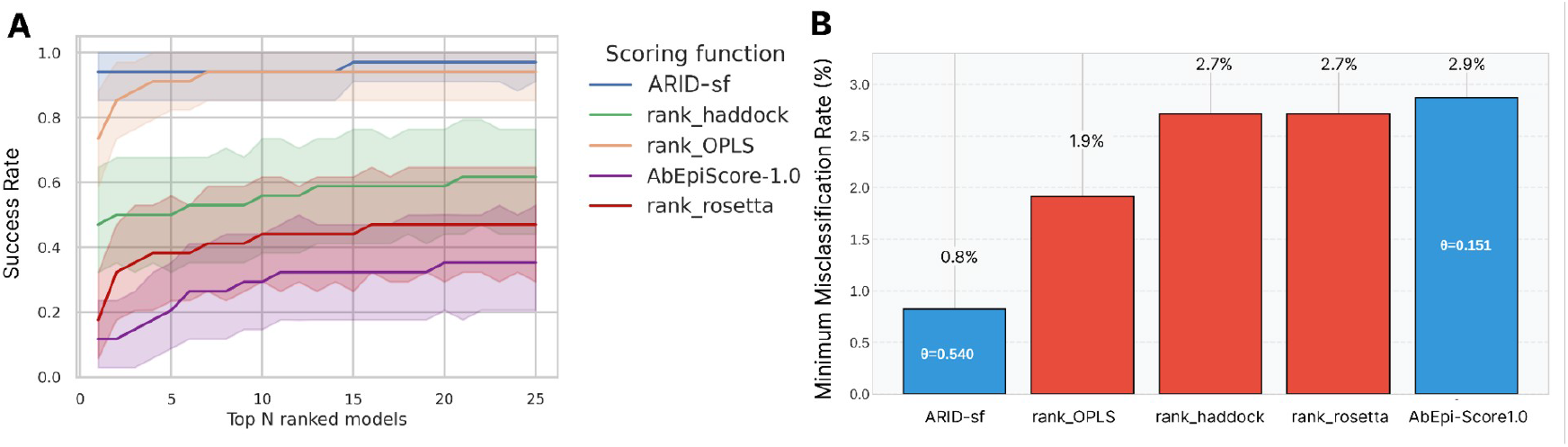
Performances of different scoring schemes for the Spd set. This experiment consists of docking Ab-Ag complexes with non native interfaces obtained via flexible refinement with HADDOCK. 12 of the 44 cases presented had an Ab modelled by AbBodyBuilder2. ***A)*** The figure shows the SR of correct (Acceptable, Medium, or High Quality) docking models (y-axis) identified within the top N ranked candidates (x-axis) using five scoring functions (ARID-sf in blue, the HADDOCK score in green, the OPLS score in orange, the Rosetta total_score in red and the AbEpiScore-1.0 in purple). ***B)*** Performances of the scoring function in classifying docking models (correct from incorrect). Minimum misclassification is evaluated using the classification threshold (displayed as Θ) that maximises the accuracy of the scoring function. A higher value signifies worse accuracy at distinguishing correct models from the rest.

In the present case, ARID-sf achieves the best result in the capacity of distinguishing high-scored decoys from correct binding modes. ARID-sf shows the lowest MCR at 1.0% (optimal threshold θ=0.540), followed by OPLS at 1.9%, while HADDOCK and AbEpiScore both show 2.5% and 2.6% (Figure 4B).

In the Spd set, the starting structures for docking were already optimized using OPLS united atom (see Methods and SM2), translating into improved performances for the rank_opls. However, the strong performance of the score comes with a caveat: its sensitivity to minor clashes increases false negative rates, potentially eliminating correct models with small steric conflicts. This highlights the advantage of machine learning approaches that can contextualize unfavorable interactions with broader interface features.

Despite the use of incorrect starting structures, ARID-sf maintains meaningful score stratification. Average predicted scores for incorrect, acceptable, medium, and high-quality models (0.07, 0.37, 0.84, and 0.94, respectively) follow the expected quality progression. While the optimal classification threshold (0.540) for identifying correct models (acceptable and higher quality) exceeds the DockQ boundary defining acceptable models (0.23), the monotonic relationship between predicted and actual quality demonstrates that ARID-sf learns and captures quality indicators.

We show in the above section that ARID-sf performs better than other scoring functions when small changes (~0.5 RMSD) are needed for the interface to reach the correct conformation. In the present section, we detail the results obtained on an even more challenging dataset, the Abd set. Indeed, Ab-Ag docking typically starts from modeled or unbound structures that sometimes deviate substantially from the bound conformations. We evaluated the scoring functions’ robustness under these conditions using the 67 unbound antibody-antigen structures from the Adb set (Zhao et al., 2024); we generated 4,000 rigid body docking models per complex using HADDOCK semi-blind protocol (see Methods and SM5,7). Of these 67 docking experiments, 39 produced at least one correct model (acceptable quality or higher). Among the 67 unbound partners in the Adb set, 11 exhibit less than 0.5Å I-RMSD with the bound complex, 25 exhibit 0.5-1Å I-RMSD, and 31 more than 1Å I-RMSD, including 4 cases with I-RMSD ranging from 2 to 2.5Å (see SM8 for metric details).

Figure 6 illustrates the scoring function SR performance overall and HR (see Methods) within the top 50 best models according to the evaluated functions. For the 39 cases, all scoring schemes show limited discrimination capability, requiring approximately 3,000 top-ranked models to achieve complete coverage (at least one identified correct model per case) (Figure 6A). ARID-sf identifies correct models within the top 50 for 17 distinct cases, compared to 9 for HADDOCK and 10 for OPLS scoring (Figure 6B).

**Figure 6:**
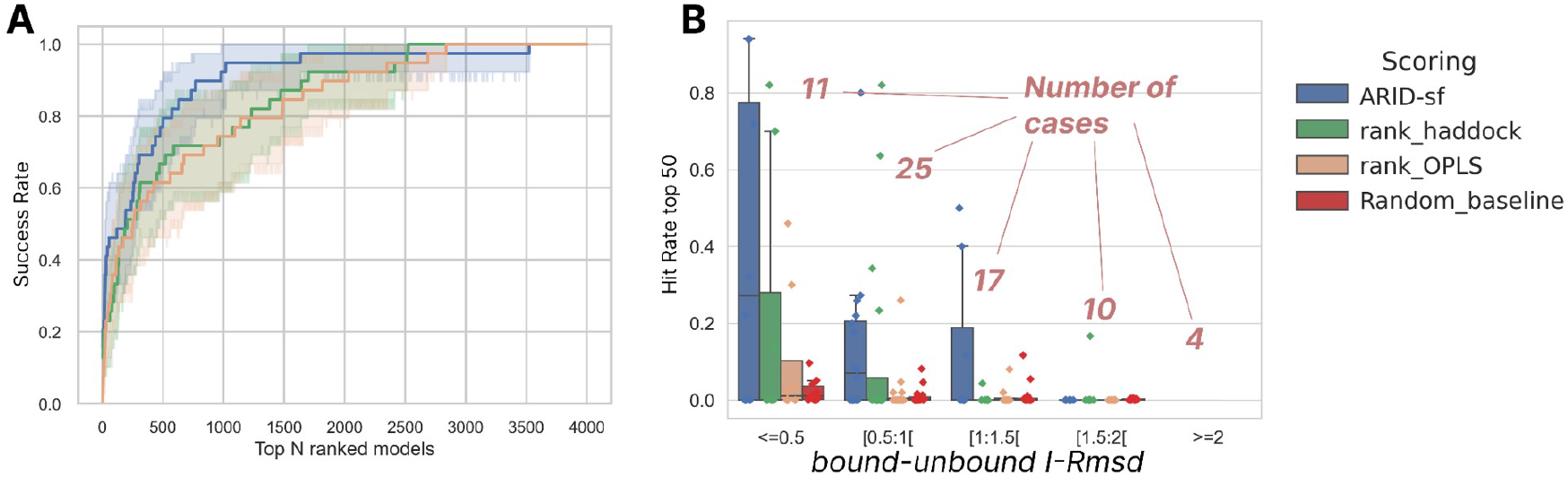
Performances of functions on the Adb set. ***A)*** SR curves showing the fraction of cases with at least one correct (Acceptable or higher quality) model in the top N selections. Shaded areas represent 95% confidence intervals. Only cases where at least one correct model has been predicted are shown. ***B)*** Effect of the increasing interface RMSD relative to the bound structures on scoring performances. The HR (different from the SR in panel *A*, see Methods) of correct models in the top 50 according to the ranking of the 3 scoring functions and random baseline (y) relative to the I-RMSD, discretized in 5 slices. Only cases where at least one correct model has been predicted have colored dots. The number of cases from each slice is indicated in beige. The random baseline is calculated by assingin random ranks to docking models.

Performance decreases strongly with the starting structures interface RMSD (I-RMSD) increase. As shown in Figure 6B, all scoring functions approach random HR performances in identifying correct poses for starting structures with I-RMSD >= 1.5 Å. However, ARID-sf outperforms the rank_haddock and rank_OPLS for cases requiring <0.51 to 1.5 Å interface RMSD to reach their bound state. The rank_haddock achieves 19% HR in the top 50 (~10 structures) for one case (Figure 6B) of the 1.5 to 2 Å interface RMSD group, highlighting the rigid-body robustness of the function for challenging cases. This also underscores the use of different weights, specifically for the term, when performing rigid-body *vs* flexible refinement docking. The separation of the repulsive and attractive terms, coupled with a rich neighboring context included in ARID-sf features, aims at identifying correct poses regardless of the docking protocol used. Although the scores and performances are different between the different docking experiments (Test, Spd, Adb), the ARID-sf achieves increased performances in each set and also on docking models produced by co-folding methods, while being the same scoring scheme throughout, getting closer to the universal scoring function goal.

Overall, ARID-sf is applicable for real case scenarios of unbound docking, when starting with incorrect structures requiring substantial changes to their interface to reach the bound state, with a noticeable enrichment in HR over the two other functions in the more challenging 0.5 - 1.5 Å interface RMSD range.

## Discussion

Evaluating docking conformations remains a fundamental challenge in computational structural biology, with Ab-Ag complexes presenting particular difficulties compared to general protein-protein interactions. The CAPRI experiment has consistently demonstrated this challenge, with Ab-Ag targets frequently classified as difficult due to the lack of performant methods for these interactions (Lensink et al., 2025). Despite advances in docking algorithms, current scoring functions struggle to identify correct poses from decoy sets, even when starting from bound conformations. This limitation has direct implications for antibody discovery pipelines, where accurate complex prediction from unbound components is critical.

Our work aims to develop ARID-sf, a machine learning-based scoring function trained on a dataset of Ab-Ag complexes that is able to rank these complexes reliably and in a reasonable amount of time, regardless of the source of the model. By testing across multiple docking scenarios (from ideal bound conformations to challenging blind predictions with modeled Abs and co-folding produced models), we provide a realistic assessment of scoring function performance. This source-agnostic function is directly applicable to HADDOCK pipelines, enhancing a docking tool already utilized for its ability to incorporate restraints in Ab-Ag studies.

## Findings

ARID-sf demonstrates improvements over physics-based scoring functions across diverse test scenarios. The performance advantage is maintained for challenging docking experiments involving structures with increasing structural perturbations compared to the bounded native state, although the raw performance decreases. Importantly, the function has learned generalizable patterns of Ab-Ag interfaces, rather than specific sequence patterns (for more details, see SM21: Influence of sequence identity on generalisation and performance).

Additionally, the function can score models generated with means other than HADDOCK, i.e, Boltz2 and AlphaFold3. Deep neural network frameworks aimed at 3D complexation inference from the amino acid sequence are becoming more and more accurate (Lensink et al., 2025). However, they still require important computational resources and time. As presented here, the generation of 25 models per case for 22 cases is already sufficient sampling to generate correct structures for about half the cases. As the evaluation of AlphaFold3 and Boltz2 for Ab-Ag complex prediction is out of the scope of this work, we will only briefly mention the accuracy in docking and focus on the scoring of both inference models. To discuss the accuracy, it would be appropriate to evaluate the redundancy between the 22 selected cases and the training set of both neural networks. It would be relevant to consider a physics-based approach for sampling the conformations, as some cases had no associated correct predictions. In ranking, both inference models are accurate, with Boltz2 slightly less precise but faster than AlphaFold3. As presented in the results, ARID-sf assigns different scores to models belonging to different structural ensembles/clusters. This is important for model selection, as it is increasingly difficult to select models if they have highly similar scores in blind experiments, and also suggests one limitation of the iPTM or ranking score from AlphaFold3.

ARID-sf does predict the quality of the Ab-Ag docking model with improved accuracy compared to the other methods presented in this work. An improvement coming from the architecture was the use of the self-attention mechanism. Initially, the function was predicting per-residue values, and the final score of the docking model was an aggregation of all its residue scores. This function weighted equally all residues of the interface, not giving more importance to critical residues. Drastic performance improvements were observed when all residues from the same structure were predicted, passing from pDockQ RMSE of 0.28 to 0.17 on the training set. Although mostly used in sequential contexts, the transformer architecture helped the model contextualize more of the ensemble of residues of a docking model for a more accurate scoring.

## Limitations

Several limitations highlight a few considerations for future development. The current training process assigns identical scores to all interface residues within a complex, potentially creating inconsistencies where locally well-formed interactions receive low scores due to global pose quality. Development of residue-specific quality metrics could address this limitation, though our experiments with residue-level LDDT scores (Mariani et al., 2013) as a predictor showed inferior performance to the current interface DockQ approach. This might be due to a need for recalibration of the set of distances used in the LDDT.

The performance gap between rigid and flexible refinement protocols highlights a challenge in scoring. Correct bound conformations possess an inherent shape complementarity that orients energy-based scoring toward native-like poses, which explains the higher performances of OPLS scoring in re-docking the Ab-Ag structures drawn from the bound structure (see Figure 3B performances in rigid re-docking). In contrast, incorrect conformations from correct starting structures are easier to identify, as their shape complementarity is low and may require substantial rearrangement to reach a local favorable molecular interface. When introducing flexibility in docking, the structural discrepancies between incorrect and correct binding poses diminish, rendering the distinction more challenging, which has been shown previously (Renaud et al., 2021; Mohseni Behbahani et al., 2022).

Training exclusively on HADDOCK-generated models may have introduced biases, particularly in the rigid orientation and refinement of the systems led by the interaction energy-based scoring function, potentially limiting generalization to models from other docking software. Although the performances were verified on Boltz2 and AlphaFold3-generated models, it is more sound to use ARID-sf in combination with HADDOCK, or at least prepare the inputs using its *topoaa* module.

Finally, ARID-sf does not take into account post-translational modifications, nor pH and ionic strength. Attempts have been made to incorporate the pH effect in the electrostatic energy calculation. A new implementation with an updated version of the force field currently used would be required for that.

## Conclusions

Elucidating the structural determinants of AbAg complexation is critical not only for understanding immune specificity but also for accelerating the development of potent vaccines and immunotherapies. One way to predict AbAg complexation is by using the new inference models like AlphaFold3 and Boltz2 to predict the complexes from the sequences alone. Yet, despite major improvements, these methods still struggle to model the 3D complexes accurately and are computationally expensive. Another way is to generate candidate conformations using physics-based docking approaches like HADDOCK3 and select the best candidates according to a scoring function. In this work, we presented ARID-sf, a scoring function specialized for Ab-Ag docking model ranking, able to accurately promote the closest to native conformation in the top ranks. Moreover, it improves on the physics-based HADDOCK3 scoring function, the Ab-Ag specialized AbEpiScore scoring function, and the Boltz2/AlphaFold3 scores for a fraction of the computational cost. Crucially, the physics-informed architecture of ARID-sf offers a distinct advantage over these end-to-end predictors: the potential for extensibility. Unlike “black-box” sequence models, our framework allows for the future incorporation of specific physicochemical conditions (such as pH effects or ionic strength), providing a path to model environmental dependencies currently inaccessible to pure deep learning approaches. In this work, we also show that the simple HADDOCK function outperforms AbEpiTope-1.0 for ranking Ab-Ag models obtained from non native docked components. The pipeline is directly usable with HADDOCK output models and is freely available at https://github.com/DSIMB/ARID-sf.git.

## Funder Information Declared

This work has been supported by the “Agence Nationale de la Recherche” [ANR-20-CE06-0029] within the frame of the Emulate project “Developing, Validating and applying computer simulation methods to Enhance the MolecULAr undersTandIng and tO eNgineer functionalized biomaterials” −EMULATE and by the “Fundação de Amparo á Pesquisa do Estado de São Paulo” [Fapesp 2020/07158–2 (F.L.B.d.S.)] and the Conselho Nacional de Desenvolvimento Científico e Tecnológico (CNPq) [307461/2025-4 (FLBdS)].

## Notes

https://github.com/DSIMB/ARID-sf.git

### Competing Interest Statement

The authors have declared no competing interest.

