## Supplementary Material for "ARID-sf: A physics-informed Deep Learning scoring function to improve Antibody-Antigen docking model ranking"

#### SM1: Clustering Parameters

**Sequence Clustering Using MMseq2.** Sequence redundancy was evaluated using MMseq2 version 14-7e284 (Steinegger and Söding, 2017) with the cluster command. All sequences from all datasets (Train/Validation/Test/Ab-blind/Ab-site) were extracted separately for antibodies and antigens into two Fasta file ensembles. The clustering was performed with a sensitivity parameter of 2 and a coverage parameter of 0. The coverage was set to 0 to prevent sequences with 100% identity from being clustered into different groups when they had different lengths. Two separate clustering runs were performed: i) for antigens, sequence identity thresholds of 60% and 20% were applied, and ii) for antibodies, sequence identity thresholds of 95% and 80% were applied.

**Structure Preprocessing Protocol.** Each case structure was processed as follows: i) antigen atoms placed first in sequential order, ii) antibody atoms placed second, and iii) all residues and atoms renumbered continuously from 1 to the total count.

**Internal Test Set Composition.** The internal test set comprises 15 cases from 15 distinct clusters, providing a proof-of-concept evaluation set on largely distinct antigen sequences and maintaining <20% sequence identity with training data Ags.

#### SM2: Detailed Composition

**Spd Set Detailed Composition.** The 100 representative cases comprised 70 targets from the Ab-blind set and 30 targets from the Ab-site set with ABodyBuilder2-modeled antibodies. Ag and Ab sequences shared less than 20% and less than 80% identity, respectively, with training sequences. Component structures (Ag and Ab) were extracted from docking models that had been subjected to flexible refinement in the Ab-Set. For each case, one docked model was randomly selected, and the Ab-Ag components were separated for subsequent re-docking. The paratope-paratope and epitope-epitope RMSD values between the selected model components and the native bound structures were calculated and fluctuated around 0.5 Å. Semi-blind HADDOCK experiments employed paratope regions as active restraints and all antigen residues with relative solvent accessibility greater than 25% as passive residues (see SM: restraint protocol). Among the 100 cases subjected to docking, 44 produced at least one acceptable to high-quality model suitable for evaluation (12 from the Ab-site modeled set, 32 from the Ab-blind set).

**Complete Case List for AlphaFold3 and Boltz2 Evaluation.** The following PDB identifiers were used for deep learning framework evaluation: '8zye', '8qh0', '8w90', '8vvl', '9njy', '8sow', '9uk5','8yub', '9jcy', '9lf8', '8wnu', '8bec', '8u08', '8jyr', '9mic', '9npi', '8be2', '9gmu', '8qf4', '8sdf',

'9oar', '9ggp'. Among these 22 cases, 16 were absent from both AlphaFold3 and Boltz2 training datasets, while 6 were absent only from AlphaFold3 training data.

**Specific Parameters for AlphaFold3 and Boltz2.** AlphaFold3 was executed with default parameters, generating 25 models per case through 5 distinct seeds with 5 predicted models generated per seed. Boltz2 was configured with the following command-line parameters: "--recycling_steps 10 --diffusion_samples 25" to replicate AlphaFold3 default prediction settings as suggested in the Boltz2 documentation.

**RMSD Calculations Between Native and Perturbed Structures.** Interface RMSD calculations for the Spd set were performed using PyMOL with the following procedure: i) extract chain A from reference structure (car) and chain B from reference structure (cbr), ii) extract chain A from model (cam) and chain B from model (cbm), iii) select epitope residues from reference (epir) as residues in car within 5 Å of cbr, iv) select epitope residues from model (epim) as residues in cam within 5 Å of cbm, v) select paratope residues from reference (parr) as residues in cbr within 5 Å of car, vi) select paratope residues from model (parm) as residues in cbm within 5 Å of cam, vii) align epir to epim using 2 cycles initially, viii) Align parr to parm using 2 cycles initially. Extract the RMSD value. Depending on the case, the number of alignment cycles was adjusted up to 5 to achieve proper alignment convergence.

**Distribution of AbEpiTarget and AbEpiScore for Spd Set Models.** Figure S1 below shows the distribution of AbEpiTarget-1.0 and AbEpiScore-1.0 scores for the Spd set docking models, stratified by quality category.

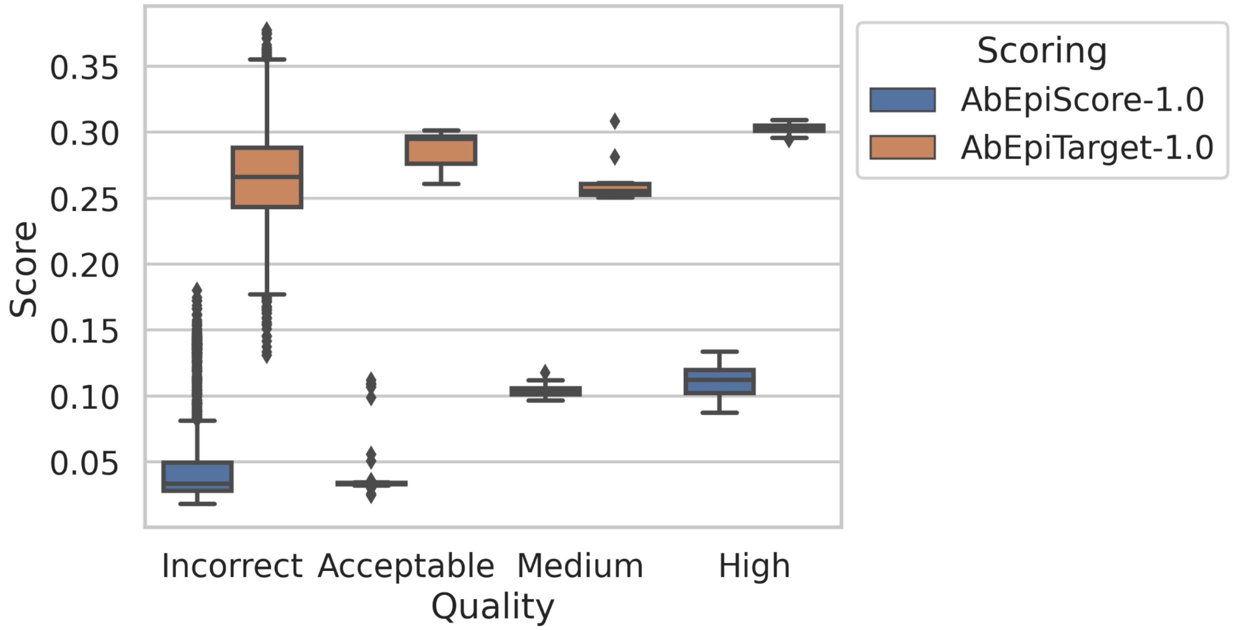


**Figure S1:** Score distributions across Incorrect, Acceptable, Medium, and High quality models

### SM3: Interface Definition

**Defining the Interface Using SASA.** Solvent accessible surface area (SASA) was calculated using Freesasa v. 2.1.2 (Mitternacht, 2016) for three configurations: i) separated antibody chain, ii) separated antigen chain, and iii) the complete complex. Residues exhibiting any change in SASA between the complex and separated states were assigned to the interface. This protocol was applied identically to both docking models and reference complexes to ensure consistent interface definitions. This protocol was only used for restraints generation using HADDOCK.

### SM4: Restraint Protocol

**HADDOCK3 Fully-Guided Protocol.** The fully-guided protocol employed for generating training models follows these steps: i) interface residues are identified from reference complexes for both Ab and antigen chains using the SASA-based method described in SM3, ii) a restraint file is created containing the complete list of interface residues for both partners, iii) both interface regions are designated as "active" residues in the ambiguous interaction restraints (AIR) framework, enabling HADDOCK to guide the docking toward the known interface region.

**HADDOCK3 Semi-Blind Protocol.** The semi-blind protocol used for the Spd and Adb sets informs on the paratope only: i) interface residues are identified from reference complexes for the Ab chain only (paratope region), ii) for the antigen, all surface-accessible residues with relative SASA greater than 25% are selected as potential interface residues, iii) a restraint file is created where paratope residues are designated as "active" while antigen surface residues are designated as "passive" in the AIR framework. This approach mimics realistic scenarios where antibody binding regions may be known while the precise epitope remains undefined.

### SM5: HADDOCK Protocol Details

**HADDOCK3 Configuration for Rigid-Body Docking.** The rigid-body docking protocol employed HADDOCK3 in local mode with the following configuration file structure:

mode = "local" molecules = [Abfile.pdb, Agfile.pdb]

[topoaa] iniseed = 766

[rigidbody] ambig_fname = "ambig.tbl" unambig_fname = "unambig.tbl" sampling = 3000

[caprieval] reference_fname = reference.pdb

The sampling parameter was set to generate 3,000 models per case for training sets and up to 4,000 models per case for certain test sets as specified in the main text. The topoaa module processes the input structures to generate HADDOCK-compatible topologies, the rigidbody module performs the docking sampling using the provided restraint files (ambig.tbl for ambiguous restraints, unambig.tbl for unambiguous restraints), and the caprieval module evaluates structural quality metrics, including DockQ and i-RMSD against the reference structure.

### SM6: Refinement Protocols

**HADDOCK3 Configuration for Flexible Refinement.** Selected rigid-body models underwent sequential refinement using the following configuration file:

mode = "local" clean = false molecules = ["ensemble.pdb"]

[topoaa] set_bfactor = false

[caprieval] reference_fname = reference.pdb

[flexref] tolerance = 10 nemsteps = 80 mdsteps_cool1 = 250 mdsteps_cool2 = 400 mdsteps_cool3 = 400 mdsteps_rigid = 250

[caprieval] reference_fname = reference.pdb

[mdref] tolerance = 10 nemsteps = 50 watercoolsteps = 50 waterheatsteps = 10 watersteps = 100

[caprieval] reference_fname = reference.pdb

### SM7: Protocol Details for Adb Set

**Semi-Blind Protocol for ABAG-Docking Benchmark.** The Adb set utilized the semi-blind protocol described in SM4, generating 4,000 rigid-body decoys per case using HADDOCK3 with the configuration specified in SM5. Interface residues for restraint generation were identified using the SASA-based method (SM3) applied to the bound reference structures provided by Zhao et al. (2024).

### SM8: Metric Details for Adb Set

**Structural Quality Metrics Calculation.** For the Adb set, structural quality assessment was performed using the caprieval module from HADDOCK3 as specified in the configuration files (SM5, SM6). The following metrics were calculated against bound reference structures: i) DockQ score for overall quality assessment and CAPRI category assignment, ii) interface RMSD (i-RMSD) measuring the positional deviation of interface residues between model and reference after optimal superposition, iii) ligand RMSD (l-RMSD) measuring the positional deviation of the entire ligand (antibody) after superposition on the receptor (antigen) [REF: Collins et al., 2024 for detailed metric definitions]. All 3D structures, bound-unbound interface RMSD values, and quality labels were obtained directly from the ABAG-docking benchmark dataset (Zhao et al., 2024).

### SM9: ESM Extraction Details

**ESM-C Model and Embedding Extraction Protocol.** The ESM Cambrian (ESM-C) model was employed to extract residue-specific representations. Embeddings were extracted at two resolution levels: i) a compact 40-dimensional vector capturing individual residue characteristics, and ii) an extended 96*3-dimensional vector encoding ag, ab, and interface-level context. These features were aggregated to reduce their lengths using mean pooling for each residue embedding, a strategy previously validated for structural and sequence feature prediction tasks (Vieira et al., 2024; Lu et al., 2025). The Ag, Ab, and Interface ESM ID (see Figure 2) were extracted using the average Ag, Ab, or interface residue embeddings (96 length). Each ESM ID has an associated token indicating the number of residues, and the Ab has an extra binary indicator for single-domain antibodies (<150 residues).

### SM10: Detailed Parameters and Validation

**Non-Bonded Interaction Parameter Specifications.** All non-bonded interaction calculations employed the OPLS united atom force field parameters in the aim of reproducing as closely as possible the HADDOCK values, as documented in the following articles (Jorgensen and Tirado-Rives, 1988; Dominguez et al., 2003; Vangone et al., 2017). The Lennard-Jones 6-12 potential was calculated using the standard functional form and explicitly decomposed into separate repulsive (r^-12 term) and attractive (r^-6 term). A switching function was applied: the switching region begins at r_on = 6.5 Å (0.65 nm) and completes at r_off = 8.5 Å (0.85 nm), with the potential forced to zero at and beyond r_off.

Coulombic electrostatic interactions were computed using the standard Coulomb equation with a shifting function applied at the same cutoff distance of r_off = 8.5 Å (0.85 nm). A distance-dependent dielectric constant of $\varepsilon = 10$ was employed. All potential calculations include contributions from heavy atoms and polar hydrogen atoms as defined by the OPLS united atom force field.

**Validation of Computed Energies Against HADDOCK Implementation.** We verified the accuracy of our potential energy calculations by systematic comparison with the OPLS potential energies computed internally by HADDOCK3. The Van der Waals and electrostatic potentials were calculated for all interface residues across diverse docking models of all quality categories.

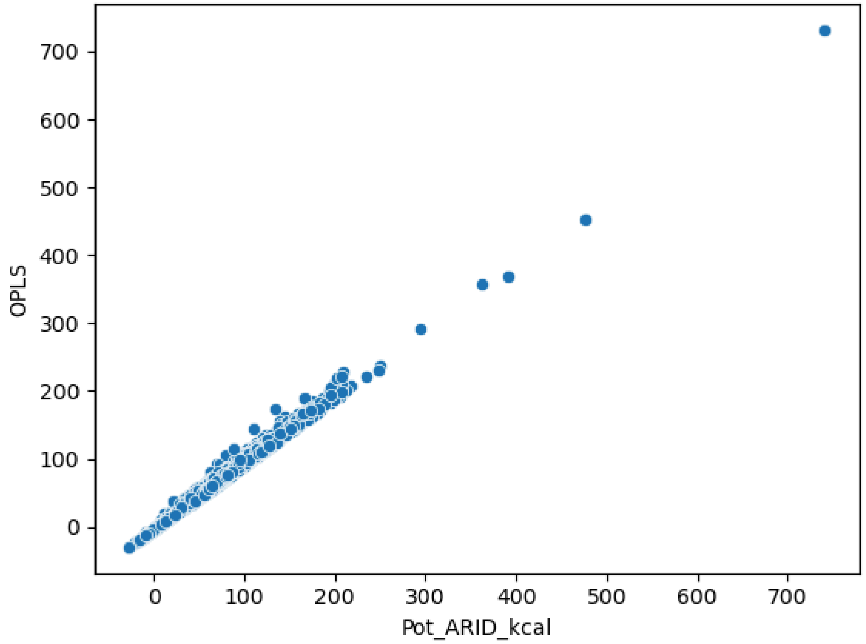


**Figure S2:** presents the correlation between ARID-calculated potentials (x-axis) and HADDOCK-reported OPLS potentials (y-axis), with both measurements expressed in kcal/mol units. The comparison reveals high correlation between the two implementations, with minor discrepancies attributed to internal optimizations of force field parameters within the HADDOCK software that are not publicly documented. These small deviations do not impact the overall fidelity of the physical interaction descriptions, as both implementations capture the same fundamental energetic trends across correct and incorrect binding poses.

### SM11: Complete Classification

**Residue Physicochemical Classification Scheme.** Interface residues were categorized into five distinct physicochemical groups based on their side-chain properties: i) Hydrophobic residues (Ala, Val, Leu, Ile, Met, Pro, Gly), ii) Polar residues (Ser, Thr, Cys, Asn, Gln), iii) Aromatic residues (Tyr, Phe, Trp, His), iv) Negatively charged residues (Asp, Glu) at pH 7, and v) Positively charged residues (Lys, Arg, protonated His) at pH 7.

### SM12: Atom Categorization Details

**Atom-Level Classification for Contact Analysis.** Individual atoms were categorized into six functional groups to capture atom-specific interaction patterns: i) Backbone atoms including the main-chain nitrogen (N), alpha carbon (Cα), carbonyl carbon (C), and carbonyl oxygen (O) atoms that define the protein scaffold, ii) Hydrophobic atoms comprising carbon atoms not classified as aromatic or backbone, usually found in aliphatic side chains, iii) Polar atoms including nitrogen (excluding backbone N), oxygen (excluding backbone O), and sulfur atoms capable of participating in hydrogen bonding or other polar interactions, iv) Aromatic atoms defined as carbon atoms within aromatic ring systems of Phe, Tyr, Trp, and His residues, and v) Charged atoms classified based on their parent residue charge state, specifically atoms within the charged moieties of Asp, Glu, Lys, Arg, and protonated His side chains. This atom-level granularity complements the residue-level classification by enabling the model to distinguish between interaction types occurring at the atomic scale, such as backbone-backbone hydrogen bonds versus side-chain electrostatic interactions.

### SM13: Detailed Calculation Method

**Voxel Grid Construction and Orientation Protocol.** For each interface residue serving as a central point, a three-dimensional voxel grid representation was constructed to capture the local spatial environment. The grid comprises a 3×3×3 array of cubic voxels, with each voxel measuring 10 Å (1.0 nm) along each edge, yielding a total sampling volume of 27,000 $Å^3$ centered on the residue. The grid orientation is established through a systematic coordinate frame definition: i) the geometric center of the grid is positioned precisely at the central residue's alpha carbon (Cα) atom coordinates, ii) the Z-axis of the grid coordinate system is defined as the unit vector pointing from the central Cα toward the geometric center of the opposite interface (paratope center if the central residue belongs to the epitope, or epitope center if the central residue belongs to the paratope), iii) the X-axis is constructed as the unit vector perpendicular to both the Z-axis and the Cα-N backbone vector (obtained via cross product, ensuring the resulting vector is orthogonal to the plane containing Z and the backbone direction), iv) the Y-axis completes the right-handed orthonormal coordinate system.

**Voxel Occupancy Feature Calculation.** Within each of the 27 voxels, atomic occupancy is quantified through volume calculation. For every atom whose center falls within a given voxel's boundaries, the van der Waals volume is calculated using element-specific radii tabulated by Bondi (1964). The total atomic volume within each voxel is computed as the sum of van der Waals volumes for all atoms it contains. This total is then converted to an occupancy ratio by dividing by the total voxel volume (1000 $Å^3$), yielding a dimensionless fraction between 0 and 1. The complementary fraction representing empty space potentially accessible to water molecules is calculated as (1 - occupancy ratio). Additional features encode the occupancy contributions separately by element type (hydrogen, carbon, nitrogen, oxygen, sulfur) and by chain identity (antibody versus antigen), enabling the model to distinguish between self-interface packing and cross-interface contacts. The complete voxel feature set for each central residue comprises: i) the 27 voxel positions defined by their (x, y, z) indices within the 3×3×3 grid, ii) total occupancy ratio per voxel, iii) element-specific occupancy ratios (C, N, O, S), iv) chain-specific occupancy ratios (Ab, Ag), and v) empty space fraction per voxel.

### SM14: Neighboring Residue Encoding

**Local Environment Description Through Nearest Neighbors.** For each interface residue designated as a central point, the local structural context is encoded through identification and characterization of the 20 nearest neighboring residues. Neighbor selection is performed based on Euclidean distances between geometric centers, defined as the mean position of all heavy atoms within each residue. Once the 20 nearest neighbors are identified for a given central residue, each neighbor is described through its own Residue features (geometric, ESM ID, and physico-chemical). The primary geometric features include: i) the center-to-center distance between the central residue and the neighbor residue, and ii) the angle defined by three points: the central residue Cα atom, the neighbor residue Cα atom, and the center of the opposite interface (paratope center for epitope residues, epitope center for paratope residues). This angle captures the relative orientation of the neighbor with respect to both the central residue and the binding interface geometry. Additional descriptors for each neighbor include its physicochemical classification (SM11), chain identity (antibody or antigen to distinguish between intra-chain and inter-chain neighbors), and ESM ID embedding vector (40) (see SM ESM-C Model and Embedding Extraction Protocol). This nearest-neighbor encoding strategy ensures that each central residue is described within its local microenvironment, enabling the neural network to learn context-dependent interaction patterns where the same residue type may contribute differently depending on its surrounding structural context.

### SM15: Neural Network Architecture and Training

**Architecture Overview.** The neural network is a transformer-based architecture designed for regression of antibody-antigen interface quality metric: the DockQ. The model processes variable-length sequences of interface residues, with each central residue represented as a 4,253-dimensional feature vector. The architecture consists of three major components: i) an input reduction and normalization layer that transforms high-dimensional features into a compact 256-dimensional representation space, ii) a stack of three transformer encoder layers that extract meaningful relationships between interfacial residues through self-attention mechanisms, and iii) a prediction head that aggregates interface-level information and generates quality predictions.

**Input Processing and Reduction.** The input reduction layer performs linear transformation from the original 4,253-dimensional feature space to a 256-dimensional space. This transformation is followed immediately by layer normalization to stabilize training and dropout regularization with probability $p = 0.4$ to prevent overfitting. The reduced representation serves dual purposes: i) it learns a more compact and informative encoding of the raw features by eliminating redundancies and emphasizing predictive patterns, and ii) importantly, it normalizes the feature scale to a suitable range for subsequent transformer processing. All feature values from the raw 4,253-dimensional input are capped to the interval [-1000, 1000] prior to reduction to prevent gradient explosion during backpropagation, with extreme values typically arising from repulsive Lennard-Jones interaction terms when atoms are in close proximity or clashing configurations.

**Positional Encoding Scheme.** To inject information about residue positions within the interface sequence, sinusoidal positional encoding is applied following the standard transformer architecture practice (Zhang et al., 2024). Since interface residues are ordered by decreasing intermolecular contact count, the positional encoding effectively captures the connectivity hierarchy within the interface, enabling the model to distinguish between highly connected core residues and more peripheral interface residues.

**Transformer Encoder Stack Specifications.** The core processing component consists of three stacked transformer encoder layers, each implementing the standard transformer encoder architecture with the following specifications: i) Multi-head self-attention mechanism with 8 attention heads (n_head = 8), where each head operates on a 32-dimensional subspace (256/8 = 32), enabling the model to attend to different aspects of the residue relationships in parallel, ii) Position-wise feed-forward network consisting of two linear transformations with a ReLU activation function between them, expanding from 256 dimensions to a hidden dimension of 1024 (4 × d_model) before projecting back to 256 dimensions, iii) Pre-normalization strategy (norm_first = True) where layer normalization is applied before the self-attention and feed-forward sub-layers iv) Residual connections around both the self-attention and feed-forward sub-layers, and v) Dropout regularization with probability p = 0.1 applied within the attention mechanism and feed-forward network to prevent overfitting.

**Attention-Based Sequence Aggregation.** Rather than using simple averaging or max pooling to aggregate the variable-length sequence of interface residues into a fixed-size representation, the architecture uses learnable attention pooling.

**Multi-Task Prediction Head Architecture.** The aggregated interface representation passes through a shared processing layer that projects from 256 dimensions to 128 dimensions using a linear transformation, followed by ReLU activation, dropout with probability p = 0.1, and layer normalization. A sigmoid activation function is applied to the outputs to constrain predictions to the [0,1] range, matching the natural scale of the target quality metrics (DockQ).

**Loss Function and Task Weighting.** Training employs a root mean square error (RMSE) loss function that computes RMSE values for the predicted metric.

**Weight Initialization Strategy.** All linear transformation layers throughout the network are initialized using Xavier uniform initialization.

**Masking Strategy for Variable-Length Sequences.** To properly handle interface sequences of varying lengths up to the maximum of 75 residues, the architecture uses masking. Padding masks are generated for each input sequence, with Boolean values indicating valid residue positions (True) versus padded positions (False). For self-attention computation, these masks prevent attention from being calculated between any residue and the padded positions by setting the corresponding attention logits to large negative values (-1×10^9) before the softmax operation, ensuring padded positions receive near-zero attention weights. Similarly, in the attention pooling mechanism, padded positions are masked using the same large negative values before the final softmax normalization, preventing them from contributing to the aggregated interface representation.

**Training Hyperparameters and Optimization.** The model was trained using the Adam optimizer with the following configuration: i) learning rate of 1×10^-5, ii) weight decay (L2 regularization) of 1×10^-4 applied to all weight parameters to prevent overfitting, iii) batch size of 1,000 interface tensors. The validation set was monitored during training to assess generalization performance. The model's weights were saved if the RMSE diminished on the validation set, epoch after epoch. The final RMSE on the validation and training set was 0.1964, and 0.1782, respectively.

### SM16: Hardware Specifications and Memory Requirements

**Training and Inference Hardware Configuration.** All model training and evaluation was performed on a computer with: i) Graphics processing unit: NVIDIA GeForce RTX 3080 with 10 GB GDDR6X memory ii) Central processing units: Dual Intel Xeon Silver 4210R processors, of 20 Cores each. iii) 64 GB or DDR4 Ram. When using the model, the amount of RAM usage can be parametrized. Typically, 10 GB of RAM is needed for extensive scoring of ~10 000 models, usage which can be splited into smaller batches using less RAM to fit the user's need.

**Performance Benchmarks and Throughput.** On the 40-CPU core configuration, the complete ARID-sf pipeline achieves throughput of 3,000 to 5,000 models per minute, with variation depending on system size (number of interface residues, total residues in complex). For a representative benchmark case consisting of 9,000 docking models of a system containing 50 interface residues, the pipeline requires approximately 10 GB of RAM and completes processing in 105.8 seconds when utilizing all 40 CPU cores for parallel feature extraction. The memory footprint scales approximately linearly with the number of models being processed simultaneously, as each model requires storage for its feature matrix (number of interface residues × 4,253 features) and intermediate calculations. The computational bottlenecks differ between pipeline stages: i) ESM-C embedding generation is GPU-bound (however, it is computed only once per system/case), ii) feature extraction (contacts, energies, voxels) is CPU-bound and benefits from parallelization, and iii) neural network inference is GPU-accelerated and relatively fast compared to feature extraction. Neural network inference automatically utilizes GPU when available and falls back to CPU computation if no compatible GPU is detected.

### SM17: Protocol-Specific Weights

**HADDOCK Score Composition Across Docking Stages.** The HADDOCK scoring function contains multiple energetic terms with weights that vary according to the docking protocol stage. The complete scoring function takes the form: HADDOCK_score = w_vdw × E_vdw + w_elec × E_elec + w_desolv × E_desolv - w_bsa × BSA, where E_vdw represents the van der Waals potential energy, E_elec represents the electrostatic potential energy, E_desolv represents an empirical desolvation energy term, and BSA represents the buried surface area at the interface. The relative weights (w_vdw, w_elec, w_desolv, w_bsa) are adjusted according to the following protocol stages:

**Rigid-Body Stage Weights.** w_vdw=0.01, w_elec=1, w_desolv=1, w_bsa=0.01

**Flexibility Refinement Stage Weights.** w_vdw=1, w_elec=1, w_desolv=1, w_bsa=0.01

**Water Refinement Stage Weights.** w_vdw=1, w_elec=0.2, w_desolv=1, w_bsa=0.00

For all evaluations in this work, the restraints penalty term was excluded from the HADDOCK score to enable fair comparison with other scoring functions that do not utilize interface knowledge. The restraints penalty quantifies violations of the user-defined distance restraints and would favor models generated with correct interface information, making cross-comparison with blind scoring functions biased.

### SM18: Detailed Processing Pipeline

**AlphaFold3 and Boltz2 Structure Processing Protocol.** Models generated by AlphaFold3 and Boltz2 deep learning frameworks required preprocessing before ARID-sf scoring to ensure topology compatibility. The complete processing pipeline follows these steps: i) Structure format conversion: The native output files from AlphaFold3 and Boltz2 in Crystallographic Information File (CIF) format were converted to Protein Data Bank (PDB) format using house scripts, ii) Topology processing: The converted PDB structures were passed through the HADDOCK3 topoaa module using a HADDOCK configuration (.cfg) file, containing the [topoaa] and [caprieval] module for metric calculations. The topology-processed structures were then scored using the standard ARID-sf pipeline with interface identification, feature extraction, and neural network inference as described in the main text and SM sections 3, 9-14, and 15.

**Confidence Score Extraction from Deep Learning Models.** For each model generated by AlphaFold3 and Boltz2, we extracted three confidence metrics provided in the output files: i) ipTM (interface predicted template modeling score), which specifically assesses the confidence in the predicted interface region and serves as a measure of how well the model's interface residues are expected to match the true structure, ii) pTM (global predicted template modeling score), which provides an overall confidence estimate for the entire predicted structure including both interface and non-interface regions, and iii) ranking_score (termed "confidence_score" in Boltz2 output files), which represents the primary metric used by each framework to rank its predicted models. As pTM consistently underperformed both ipTM and ranking_score for the task of identifying correct docking poses versus incorrect poses, all comparative evaluations focus on ipTM and ranking_score as the representative confidence measures from these deep learning frameworks.

**Rosetta Score Calculation.** Rosetta Interface Analyzer was applied to each model in the Spd set using the REF2015 scoring function with default parameters. The total_score output from the Interface Analyzer, representing the overall energetic favorability of the interface, was extracted and used as the ranking score for comparison with ARID-sf and other scoring functions.

#### SM19: Results Complete Test Set

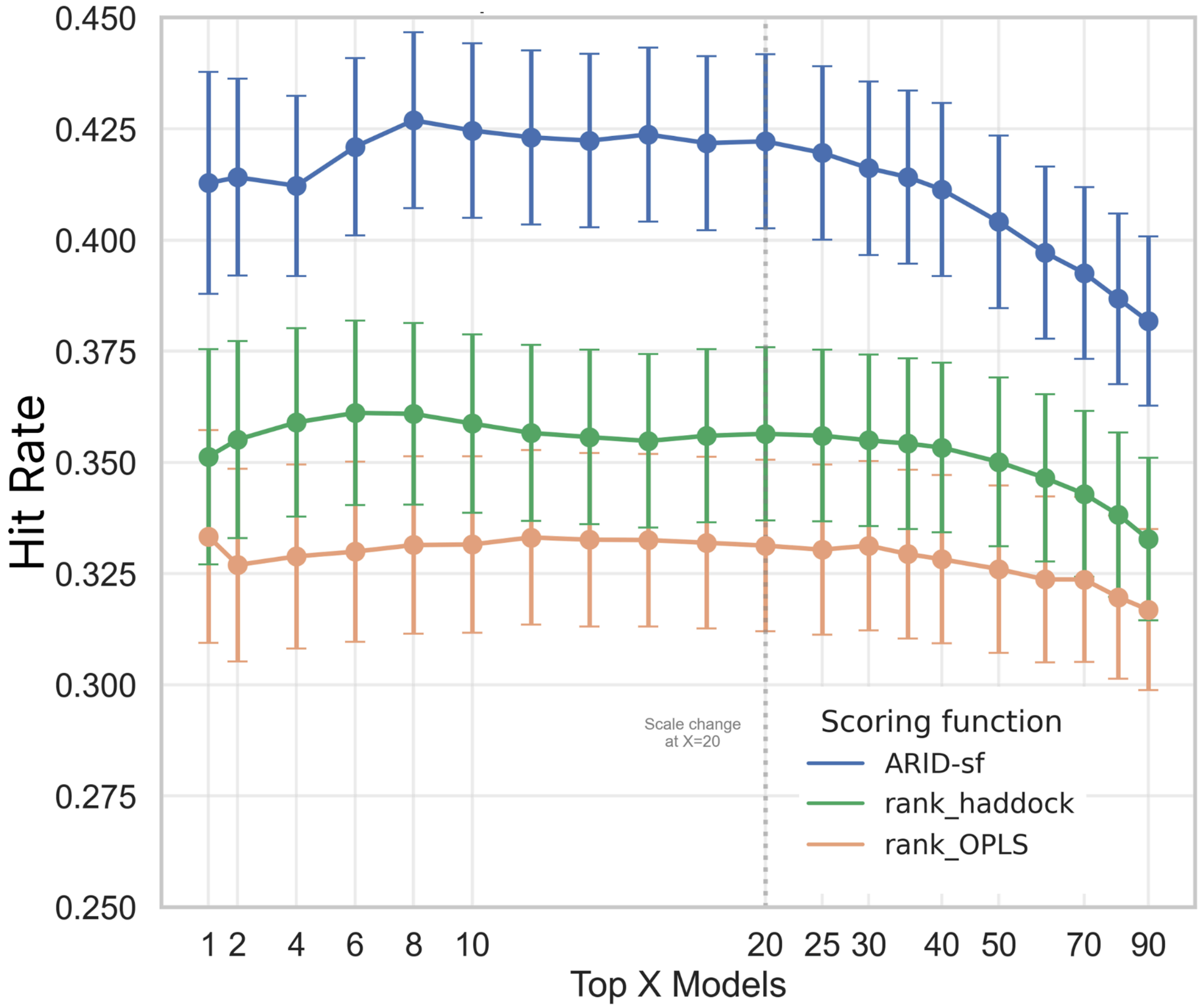


**Figure S3**: Hit rate curves showing the fraction of correct (acceptable quality and up) models within the top X ranked candidates for ARID-sf (blue), HADDOCK score (green), and OPLS score (orange). A value of 0.4 at Top 10 indicates that 40% of the ten highest-ranked models are correct.

Hit rate analysis reveals the expected proportion of correct models at each ranking depth (Figure S3A), highlighting the scoring function's ability to correctly order the top predictions first. ARID-sf maintains greater performance: 41.2% and 42.4% of correct models are ranked in the top 1 and 10, respectively, against 35.1% and 35.8% for HADDOCK and 33.3% and 33.1% for OPLS. The performances of the functions are not significantly different when considering only the Ab-site subset of cases, with 54.1% +/-39.2, 51.6% +/-42.8, and 51.1% +/-42.9 for ARID-sf, HADDOCK, and OPLS, respectively. This is due both to the nature of the set, with modelled Ab representing a more realistic scenario with model quality having direct implications on docking success (Harmalkar *et al.*, 2025), and the amount of sampling, which is only 250 per case (a low number considering HADDOCK guidelines for Ab-Ag docking).

### SM20: Results Spd set

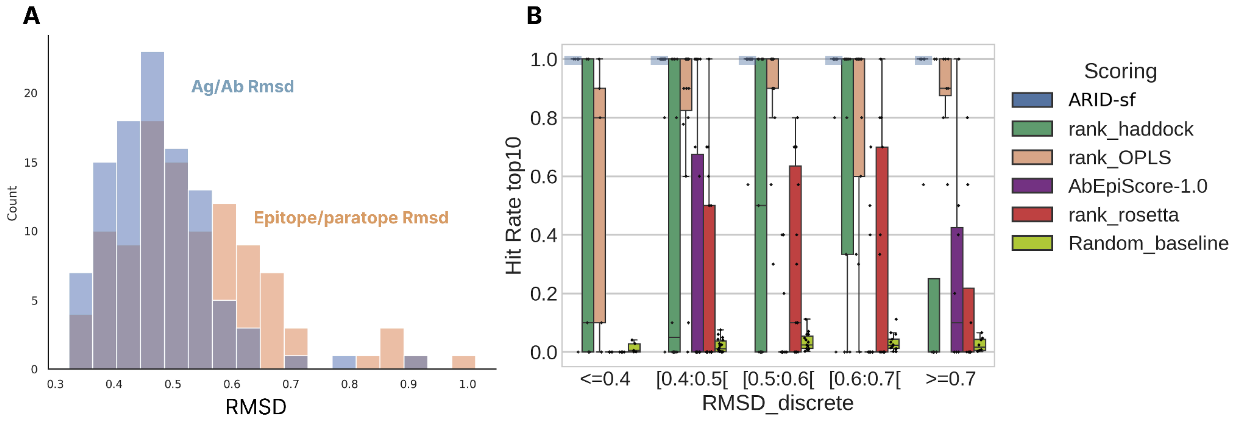


**Figure S4**: A) RMSD distribution for the Ab and Ag structures used for docking relative to the native Ab and Ag structures. The RMSD for the epitope and paratope is shown in orange, and the global RMSD is shown in blue. Each value (Ag, Ab, paratope, epitope) is represented. B) Effect of the increasing epitope/paratope RMSD relative to the native structures on scoring performances. The Hit rate of correct models in the top 10 according to the ranking of the 4 scoring functions (y) relative to the epitope/paratope RMSD, discretized in 5 slices.

### SM21: Influence of sequence identity on generalisation and performance

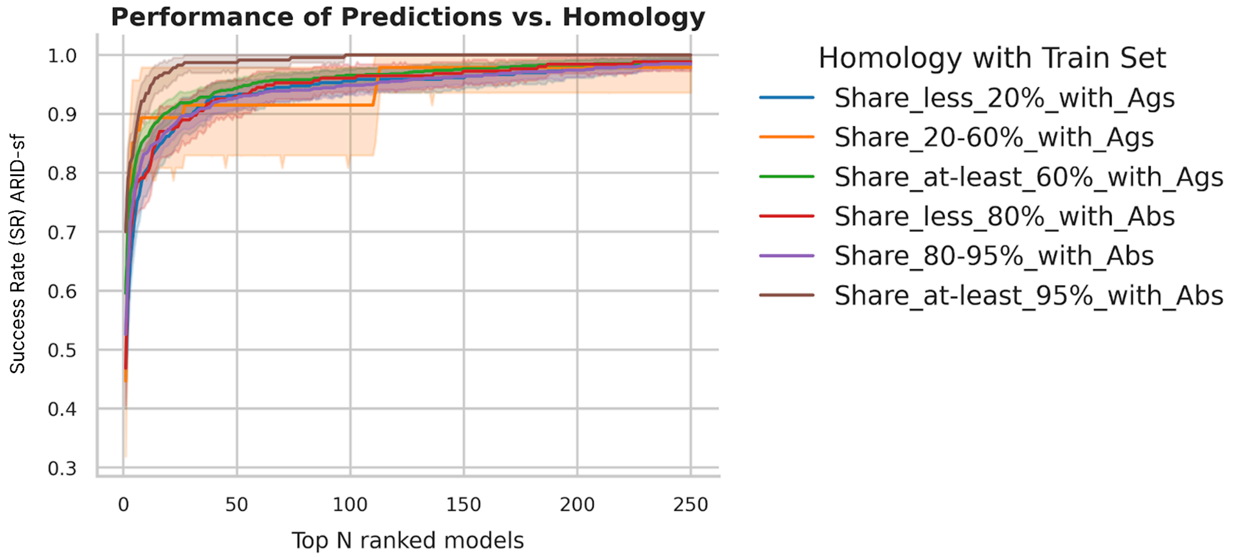


**Figure S5**: Performances for different groups of cases based on sequence identity with the set of cases used to train ARID-sf. The figure shows the fraction of cases (y-axis) where at least one correct model has been ranked in the top N (x-axis). The different colored lines represent the groups of cases based on their Ag or Ab sequence identity with the Train set. The analysis encompasses 1220 total cases distributed as follows: 362 cases sharing <20% identity with training Ags, 47 cases with 20-60% antigen identity, 811 cases with ≥60% antigen identity, 254 cases sharing <80% Ab identity, 736 cases with 80-95% Ab identity, and 230 cases with ≥95% Ab identity. The semi-transparent areas represent 95% confidence intervals.

To evaluate whether ARID-sf generalizes beyond its training data or simply memorizes sequence interface patterns, we analyzed performance across different sequence identity ranges. Cases gathered from all different sets used in this work were grouped using the same MMseq2 clustering thresholds employed for dataset partitioning: 20% and 60% for Ags, 80% and 95% for Abs. Cases that appear twice in Ab-blind and Ab-site are grouped and counted once.

Figure S5 demonstrates consistent performance across sequence identity ranges except for cases sharing at least 95% Ab identity, which displays higher performance. Success rates vary by less than 0.1 across most groups, with overlapping confidence intervals indicating no statistically significant differences. Yet, there is still a progression between distant sequences (<20% Ag or <80% Ab identity) towards more similar sequences (>60% Ag or >95% identity). This result highlights that the function has probably memorized some sequence patterns, and performs slightly better when sequences are closer to its training set. The group of cases sharing 95% or more Ab sequence identity has higher than average performances compared to the other groups, suggesting that the neural network might have learned to predict more accurately Ab that are similar to its training set. It is important to note that this group contains various levels of identity in the Ag side, with 226 distinct groups of sequences sharing less than

20% identity. This result suggests that Ab diversity plays a more important role than expected in Train/Validation/Test splits.

The comparable performance across sequence identity ranges provides evidence that ARID-sf learns generalizable physicochemical and structural principles rather than specific sequence patterns. This generalization capability is important for practical applications, where novel Ab-Ag pairs share limited sequence identity with existing complexes in structural databases. The consistent performance suggests ARID-sf will maintain reliability even for entirely novel therapeutic Ab candidates or emerging antigenic targets.
